# Genome edited colorectal cancer organoid models reveal distinct microRNA activity patterns across different mutation profiles

**DOI:** 10.1101/2021.12.13.472432

**Authors:** Jonathan W. Villanueva, Lawrence Kwong, Teng Han, Salvador Alonso Martinez, Michael T. Shanahan, Matt Kanke, Lukas E. Dow, Charles G. Danko, Praveen Sethupathy

## Abstract

Somatic mutations drive colorectal cancer (CRC) by disrupting gene regulatory mechanisms. Distinct combinations of mutations can result in unique changes to regulatory mechanisms leading to variability in the efficacy of therapeutics. MicroRNAs are important regulators of gene expression, and their activity can be altered by oncogenic mutations. However, it is unknown how distinct combinations of CRC-risk mutations differentially affect microRNAs. Here, using genetically-modified mouse intestinal organoid (enteroid) models, we identify 12 different modules of microRNA expression patterns across distinct combinations of mutations common in CRC. We also show that miR-24-3p is aberrantly upregulated in genetically-modified mouse enteroids irrespective of mutational context. Furthermore, we identify an enrichment of miR-24-3p predicted targets in downregulated gene lists from various mutational contexts compared to WT. In follow-up experiments, we demonstrate that miR-24-3p promotes CRC cell survival in multiple cell contexts. Our novel characterization of genotype-specific patterns of miRNA expression offer insight into the mechanisms that drive inter-tumor heterogeneity and highlight candidate microRNA therapeutic targets for the advancement of precision medicine for CRC.

## Introduction

Colorectal cancer (CRC) is estimated to be the third most diagnosed cancer and the second leading cause of cancer-related death worldwide^1^. A major challenge in treating CRC patients is that molecular differences across patients’ tumors, or inter-tumor heterogeneity, can lead to highly variable patient outcomes^2-4^. Recent advances in the understanding of CRC inter-tumor heterogeneity have led to substantial improvements in the therapeutic strategies utilized to treat CRC patients^4-6^ One notable example is how tumors are screened for *KRAS, NRAS*, and *BRAF* mutations to determine eligibility for anti-EGFR monoclonal antibody treatment^5,6^. This example represents only the beginning of the promise of personalized approaches for CRC, and strongly motivates the goal of understanding how different combinations of somatic mutations in key oncogenes and tumor suppressors promote molecular variability across tumors.

Mutation status plays a key role in inter-tumor heterogeneity through genotype-specific alterations of gene regulatory mechanisms that control tumor growth and development^7-9^. Unique combinations of driver mutations have been shown to lead to novel cancer phenotypes, including resistance to WNT inhibitors in intestinal mouse models of CRC^10-12^. However, most studies that investigate the effects of genetic alterations on gene regulatory mechanisms focus on the effects of individual mutations^8,13,14^. Therefore, there is a critical need to investigate how combinations of distinct CRC mutations alter regulatory mechanisms and drive novel cancer phenotypes.

MicroRNAs (miRNAs) are small, ∼22 nt non-coding RNAs that canonically function as post-transcriptional, negative regulators of gene expression. It has been well documented that abnormal activity of certain miRNAs can initiate and/or exacerbate disease phenotypes, including cancer^15-17^. Although there remain some challenges to miRNA-based therapeutics (as with many other classes of molecular therapy), several have shown promise in pre-clinical models of cancer (such as miR-10b in breast cancer^18^ and glioblastoma^19^) and some have been nominated for clinical trials^20^ and/or are currently in different phases of clinical trials^21^. Numerous studies have demonstrated that miRNAs are significantly altered in CRC tissues^22-24^. While miRNA-based therapies have been proposed for CRC^25^, to our knowledge none are currently in clinical trials. Moreover, importantly, it remains unknown how different combinations of driver mutations affect miRNA profiles and how this promotes unique tumor phenotypes.

A major challenge in evaluating how combinations of mutations affect miRNA profiles has been a lack of appropriate cellular models. Primary tumors harbor tens to hundreds of non-silent mutations and are therefore not ideal for evaluating the effects of specific genotypes^2^. Additionally, primary tumors are highly heterogenous and this limits our ability to assess mutation-specific miRNA alterations in the epithelium where CRC tumors form. CRC cell models also have several mutations^26^ and are limited in their ability to recapitulate the biology of the intestinal epithelium. To address these limitations, researchers have developed genetically modified organoid models that mimic the physiology of the intestinal epithelium. Using gene editing tools (CRISPR/Cas9, Cre), specific combinations of mutations can be induced to evaluate their impact on cell behavior and/or sensitivity to therapeutics^12,27^. To our knowledge, these state-of-the-art intestinal model systems have not yet been used to study mutation-specific changes to miRNA profiles.

To address the important knowledge gaps mentioned above, we leverage genetically modified mouse small intestinal epithelial organoids (termed enteroids) to characterize how miRNA profiles change in response to different combinations of CRC driver mutations. Using small RNA-seq, we define different patterns of miRNA expression across various genotypes. In doing so, we highlight the dominant role of Tgf-B signaling in the regulation of predicted tumor suppressor miRNA, miR-375-3p. By leveraging this mouse enteroid data, in conjunction with small RNA-seq data from human primary colon tumor data from The Cancer Genome Atlas (TCGA)^2^, we find that miR-24-3p is up-regulated across all mutational contexts. Additionally, we observe an enrichment for predicted miR-24-3p targets in genes downregulated in multiple CRC contexts. Additional studies in multiple cell models demonstrate that miR-24-3p inhibition results in a significant decrease in cell viability by inducing apoptosis. Finally, we perform integrative analysis of RNA-seq and chromatin run-on sequencing (ChRO-seq)^28^ to identify *HMOX1* and *PRSS8* as genes subject to strong post-transcriptional regulation by miR-24-3p in CRC. Overall, this study offers, to our knowledge, the first genome-scale characterization of miRNA patterns across distinct combinations of CRC driver mutations, provides new insight into the molecular mechanisms that drive inter-tumor heterogeneity, and defines candidate miRNA targets for future therapeutic development in CRC.

## Results

### Genetically modified enteroids exhibit mutation-specific variation in miRNA expression

To characterize the effect of genotype on miRNA expression we performed small RNA-seq on mouse enteroids that harbor different combinations of CRC mutations (**Figure 1A, Table S1**). We focused on mutations in genes that are in signaling pathways commonly dysregulated in CRC according to The Cancer Genome Atlas (TCGA)^2,12,29-31^: Wnt (*Ctnnb1, Apc*, and *Rspo3*; 181/195 tumors in TCGA contain at least one mutation affecting this pathway), p53 (*p53*; 120/195 tumors in TCGA contain at least one mutation affecting this pathway), Mapk (*Kras*; 122/195 tumors in TCGA contain at least one mutation affecting this pathway), and Tgf-B (*Smad4*; 70/195 tumors in TCGA contain at least one mutation affecting this pathway). Using miRquant 2.0, a small RNA-seq analysis tool^32^, we profiled miRNAs across enteroids with 9 different genotypes. Principal component analysis (PCA) revealed that miRNA profiles stratify enteroid samples by mutational combinations (**Figure 1B**). Moreover, the majority of mutant enteroids are clearly separated from wild-type (WT) in the PCA plot. The analysis also shows that *Rspo3* mutant enteroids are most similar to WT controls, which is in line with previous morphological and RNA-seq comparisons^31^. Therefore, *Rspo3* mutants were not incorporated into the downstream analyses.

**Figure 1:**
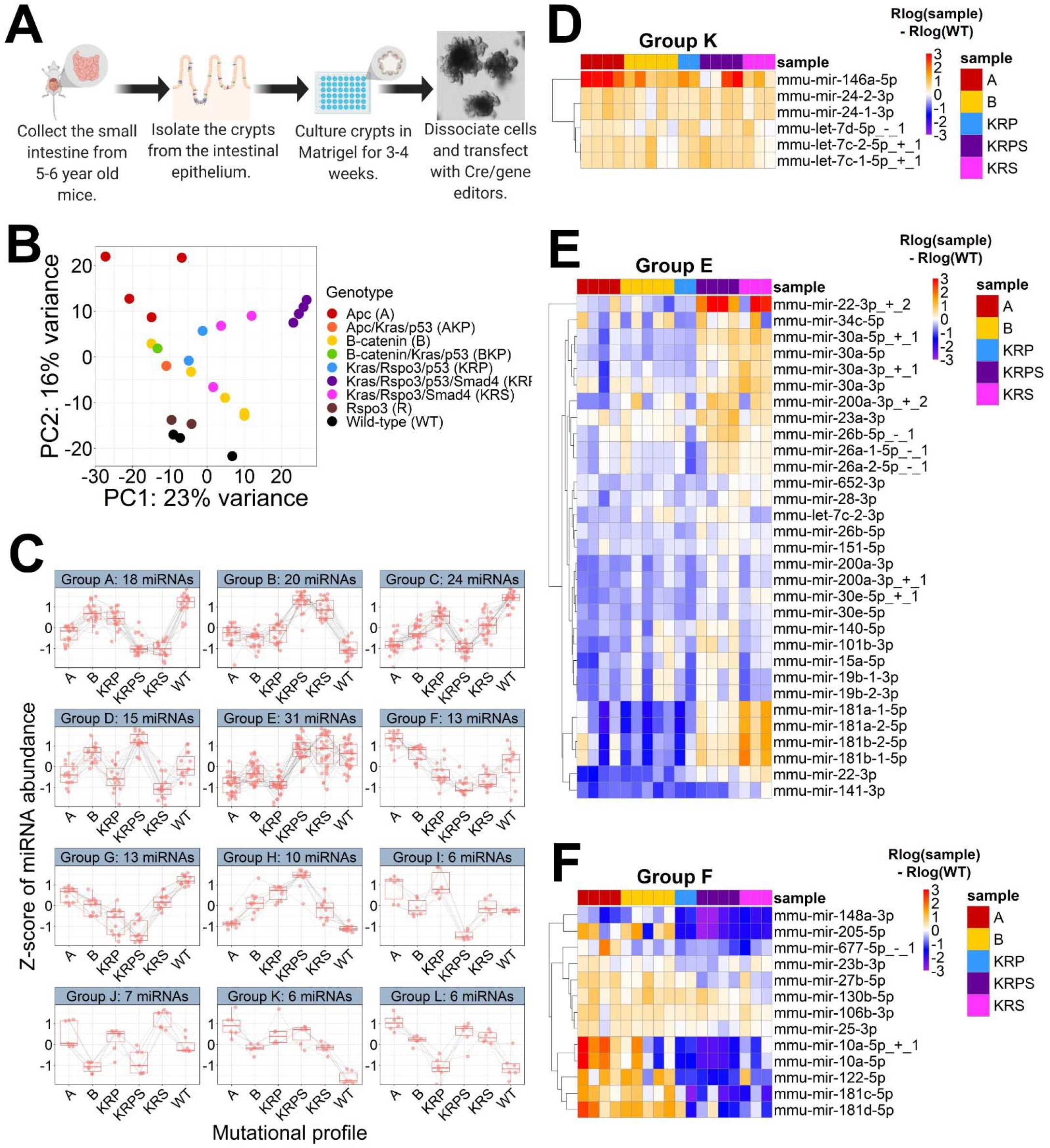
Genetically modified enteroids exhibit mutation-specific variation in miRNA expression. (**A**) Diagram illustrating how enteroid models were generated (Created with BioRender.com). (**B**) Principal component analysis (PCA) plot generated using miRNA expression profiles from *Apc* (A; n=4), *Apc*/*Kras*/*p53* (AKP; n=1), *Ctnnb1* (B; n=5), *Ctnnb1*/*Kras*/*p53* (BKP; n=1), *Kras*/*Rspo3*/*p53* (KRP; n=2), *Kras*/*Rspo3*/*p53*/*Smad4* (KRPS; n=4), *Kras*/*Rspo3*/*Smad4* (KRS; n=3), *Rspo3* (R; n=2) mutant enteroids, and wild-type (WT; n=3) controls. (**C**) Z-score of miRNA abundance for the 12 modules of miRNA expression, each with greater than 5 miRNAs in the module, as defined by DEGReport. Only miRNAs with baseMean > 500 and p-adj < 0.05 following DESeq2 likelihood ratio test (LRT) were included in the analysis. (**D-F**) Heatmaps show the magnitude of change in miRNA expression relative to WT by subtracting rlog normalized miRNA expression for each enteroid sample by the rlog average WT expression. Heatmaps shown are for Group K, Group E and Group F as defined by DEGReport. Color intensity shows the difference between rlog normalized miRNA expression and average WT. Color scale minimum saturates at -3 and maximum saturates at 3.

Next we sought to define miRNA expression patterns across the 6 genotypes for which we have at least two biological replicates. Specifically, we performed a likelihood ratio test using DESeq2, which revealed 175 miRNAs with significant expression variation across genotypes (p-adj<0.05, baseMean>500). We grouped these miRNAs into 12 distinct expression profiles, or “modules”, using DEGreport^33^ (**Figure 1C, Supplemental Figure 1**). Group K (**Figure 1D**) is composed of miRNAs that exhibit a similar increase in expression across all genotypes relative to WT. One prominent example of a Group K miRNA is miR-146a-5p^34,35^, which functions as an oncogenic miRNA in CRC. All remaining modules exhibits non-uniform effects on miRNA expression; that is, larger changes in specific genotypes compared to others.

We observe multiple miRNAs, such as miR-10b-5p and miR-374-5p, that exhibit uniquely aberrant expression in *Kras*/*Rspo3*/*p53*/*Smad4* (KRPS) mutant enteroids, which possess the greatest mutational burden (**Supplemental Figure 1**}). These miRNAs may highlight a potential mechanism by which the combination of *KRAS, P53*, and *SMAD4* mutations promotes particularly severe patient outcomes^36,37^. However, we also observe miRNA modules that display the largest expression change in enteroids with the lowest number of mutations. One such example is Group E (**Figure 1E**), in which miRNAs change the most relative to WT in *Apc* (A), *Ctnnb1* (B), and *Kras*/*Rspo3*/*p53* (KRP) mutant enteroids. This group includes tumor suppressor miRNAs such as miR-30a-5p^38,39^ and miR-141-3p^40,41^. Although KRPS mutant enteroids harbor the largest number of mutations, the miRNAs in Group E exhibit only a slight elevation in this genotype. Taken together, this data supports the conclusion that the observed changes in miRNA expression are associated with specific mutational contexts, and not just a result of total mutation burden.

Some modules, such as Group F (**Figure 1F**), clearly highlight miRNAs associated with a particular pathway. MiRNAs in this group are elevated in mouse enteroids with either A or B mutant genotypes, in which we expect the strongest perturbation of the Wnt pathway. Multiple of these miRNAs, such as miR-10a-5p^42^ and miR-181d-5p^43^, have been shown to be responsive to alterations in Wnt signaling. Additionally, this group contains miRNAs, such as miR-181c-5p^44^ and miR-181d-5p^45^, that are associated with more severe CRC phenotypes. This data provides valuable insight into the role of aberrant signaling pathways on miRNA expression in the intestinal epithelium under different mutational contexts.

### Modification of Tgf-B/Smad4 signaling is sufficient to drive miR-375-3p expression in mouse enteroids

To further explore how mutations in one specific pathway can play a prominent role in the expression of miRNAs, we turned to modules in which the most significant changes in miRNA expression occur in enteroids harboring a *Smad4* mutation. Group B consists of miRNAs that exhibit the highest expression in the enteroids with *Kras*/*Rspo3*/*Smad4* (KRS) and KRPS genotypes (**Figure 2A**), whereas Group A consists of miRNAs with the lowest expression in these two genotypes (**Figure 2B**). The latter includes miR-375-3p (**Figure 2C**), which has been reported to function as a tumor suppressor in several different cancer types^46-48^.

**Figure 2:**
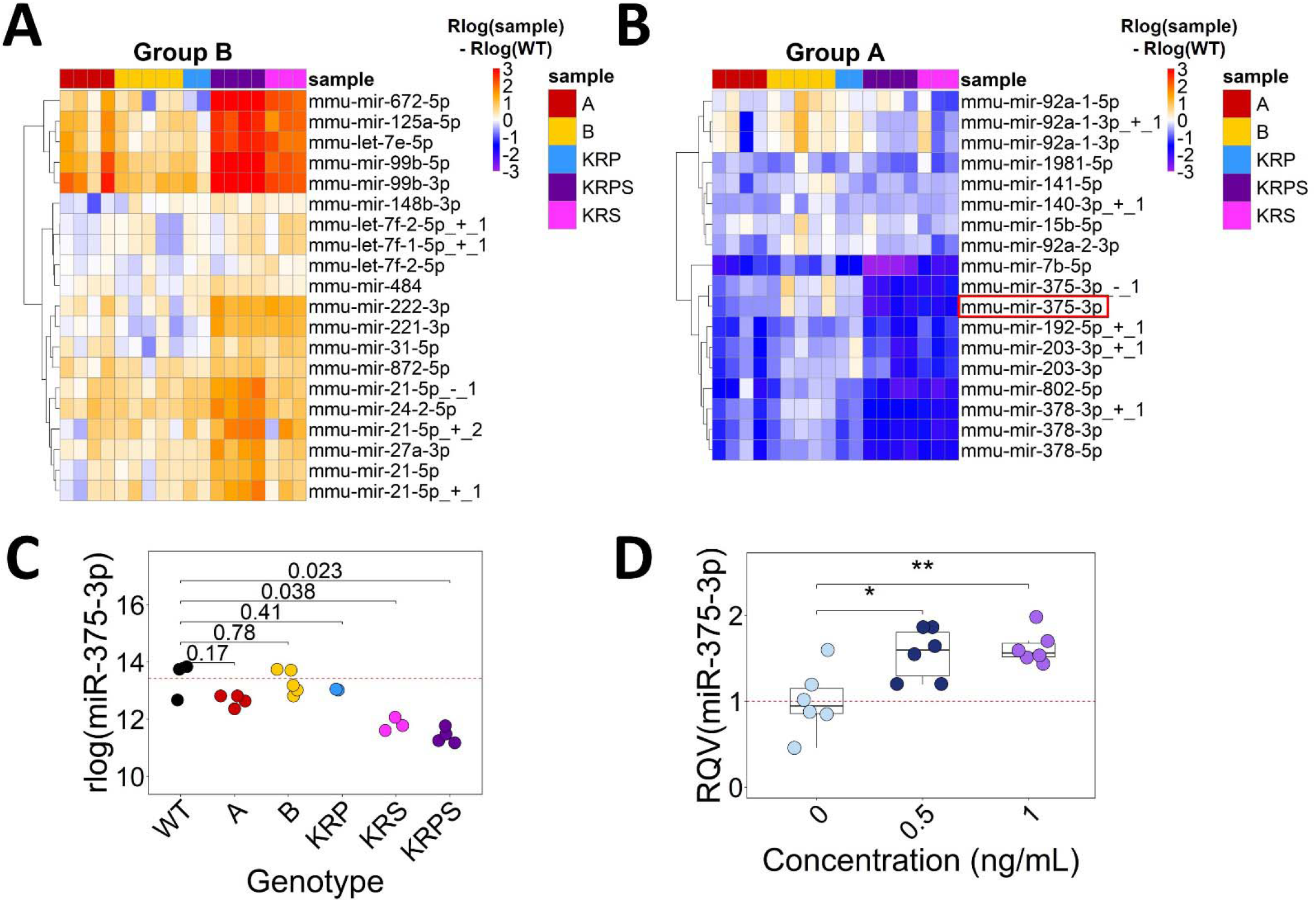
Smad4 signaling is a major driver of miR-375-3p expression in mouse enteroids. (**A, B**) Heatmaps show the magnitude of change in miRNA expression relative to WT by subtracting rlog normalized miRNA expression for each enteroid sample by the rlog average WT expression. Heatmaps shown are for Group B and Group A as defined by DEGReport. Color intensity shows rlog normalized miRNA expression in each genetically modified enteroid sample subtracted from average WT. Color scale minimum saturates at -3 and maximum saturates at 3. (**C**) Normalized miR-375-3p expression from small RNA-seq in each genotype. (**D**) MiR-375-3p expression from RT-qPCR following 0, 0.5, or 1 ng/mL treatment of mouse enteroids with recombinant human TGF-B1. Significance in (**C**) and (**D**) determined according to two-tailed Welch t-test. *p<0.05, **p<0.01, ***p<0.001.

The only difference between KRP and KRPS is the presence of the *Smad4* knockout mutation. Our findings in **Figure 2C** suggest that the loss of *Smad4* has a prominent suppressive effect on miR-375-3p, which directly motivates the hypothesis that Tgf-B signaling is sufficient to increase miR-375-3p expression in mouse enteroids. To test this hypothesis, enteroids from WT B62J mice were treated with 0, 0.5, or 1 ng/mL TGF-B1 for 3 days and changes in miR-375-3p expression were quantified using RT-qPCR. Cultures treated with TGF-B1 exhibit an expected decrease in enteroid number and elevated expression of Tgf-B regulated genes (**Supplementary Figure 2**)^49^. TGF-B1 treatment also results in a significant increase in miR-375-3p compared to control (**Figure 2D**). These results confirm our hypothesis that the candidate tumor suppressor miRNA, miR-375-3p, from Group A is most strongly driven by changes in Tgf-B/Smad4 signaling in the intestine.

### Identification of differentially expressed miRNA regulators of gene expression across various genetically modified mouse enteroid models

We next investigated miRNAs that are broadly differentially expressed across mutational contexts. These miRNAs may regulate CRC phenotypes across a broad range of genotypes and therefore could represent attractive candidates for generalized therapy. Using miRbase, we filtered for miRNA strands that are most frequently incorporated into the RNA-induced silencing complex (termed guide miRNAs). We identify 19 guide miRNAs that are significantly differentially expressed when comparing mutant enteroids to WT control (**Figure 3A, B**; DESeq2^50^ baseMean >500, >1.5x fold change, p-adj<0.05). We next performed pair-wise comparisons between each mutant genotype (with n>1) and WT controls and found 10 miRNAs that exhibit consistent up- or downregulation (DESeq2 fold change >1.5x) across all five comparisons (**Figure 3C, D**).

**Figure 3:**
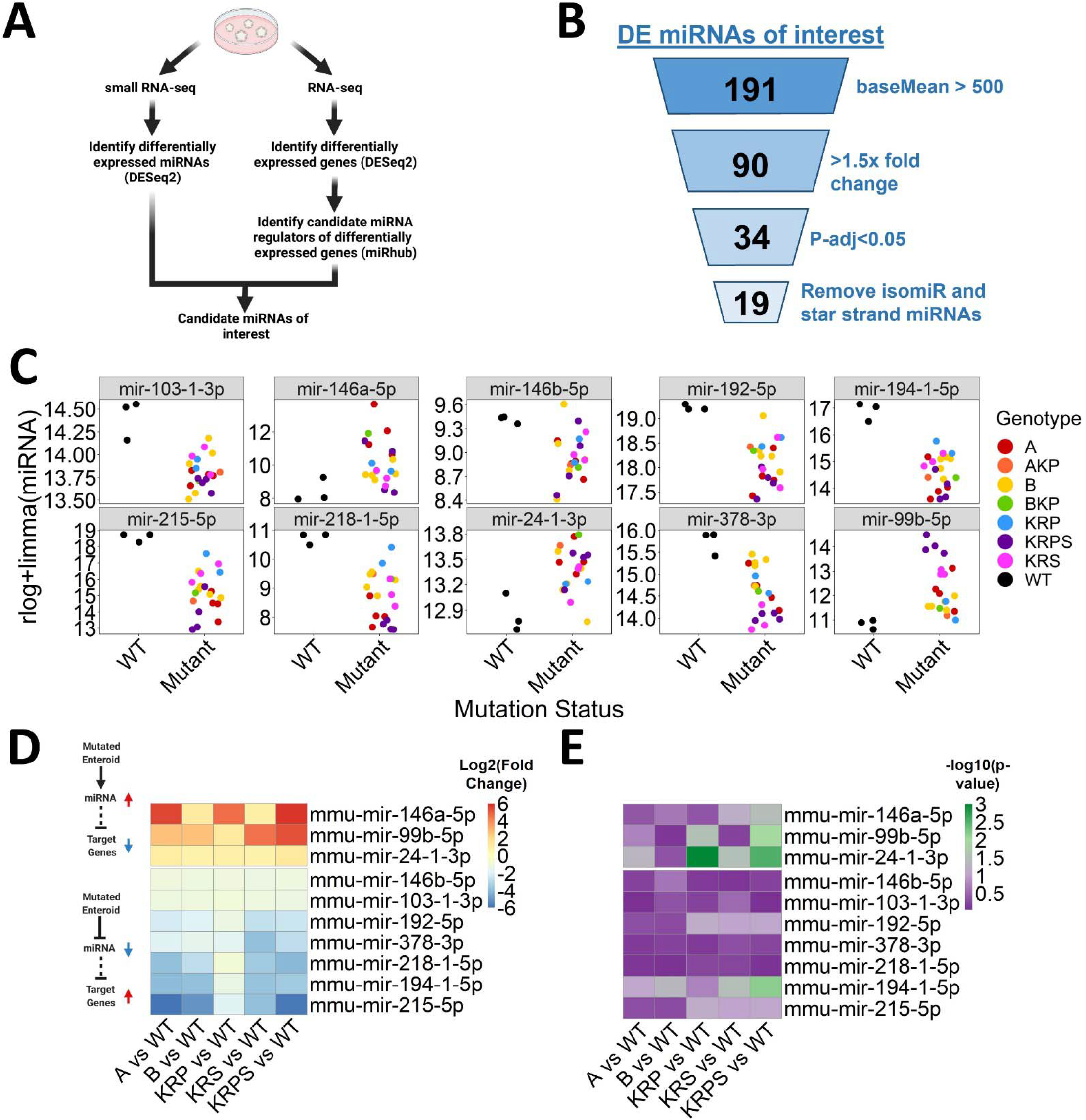
Identification of differentially expressed miRNA regulators of gene expression across genetically modified mouse enteroid models. (**A**) Schematic of the strategy utilized to identify miRNAs differentially expressed across a broad range of genotypes (Created with BioRender.com). (**B**) Results of the strategy highlights 19 guide miRNAs that are significantly differentially expressed (DESeq2 p-adj < 0.05, baseMean>500, >1.5x fold change) in mutant genotypes relative to WT. (**C**) As shown by the rlog normalized counts, 10/19 miRNAs highlighed in (**B**) are differentially expressed in the same direction when comparing mutant genotypes (with n>1) to WT. In the case of miRNAs for which both paralogs were identified as differentially expressed, only one paralog is shown. (**D**) Heatmap showing log2 fold change for miRNAs shown in (**C**). Color intensity represents the log2 fold change relative to WT. (**E**) Heatmap showing -log10(p-value) of target site enrichment, calculated by miRhub (cons1) for each differentially expressed miRNA from (**D**), in the list of genes that are differentially expressed (DESeq2 p-adj < 0.05, baseMean>500, >1.5x fold change) in the opposite direction of the miRNA.

Given that changes in miRNA expression don’t necessarily correlate with changes in activity, we next performed RNA-seq in the same mutant enteroid models (**Table S2, Supplementary Figure 3**) to evaluate gene expression changes in predicted targets of the most altered miRNAs. Using differentially expressed genes from each genotype (compared to WT) as input for our previously described statistical simulation tool, miRhub^51^, we can narrow down candidate miRNA regulators of gene expression changes across mutational contexts. MiRhub analysis identifies one upregulated miRNA with a significant enrichment (**Figure 3E**; p-value<0.05 in at least 3 out of 5 WT vs mutant enteroid comparisons) of predicted gene targets in the lists of downregulated genes. From the downregulated miRNAs, miRhub highlights one miRNA with a significant enrichment of predicted gene targets in the lists of upregulated genes (**Figure 3E**; p-value<0.05 in at least 3 out of 5 WT vs mutant enteroid comparisons). We highlight these two miRNAs, miR-24-3p and miR-194-5p, as candidate regulators of gene expression across various mutational contexts.

### miR-24-3p is a candidate regulator of gene expression and cancer phenotypes in the human colon

To place our mouse enteroid studies in a more clinically relevant context, we downloaded small RNA- and RNA-seq data from human primary colon adenocarcinoma and non-tumor tissue analyzed by TCGA^2^. After removing miRNAs with average expression under 1000 reads per million mapped to miRNAs (RPMMM) in either the tumor or non-tumor condition, we find 65 miRNAs with a significant change of expression in the tumor compared to non-tumor control (**Figure 4A**; fold change >1.5x, p-adj<0.05). Next, we identify 3190 differentially expressed genes (DESeq2; average expression > 1000 normalized counts, >1.5x fold change, p-adj<0.05). Of the 65 miRNAs that are altered in human CRC tumors, 17 exhibit a significant enrichment of predicted targets among genes that change significantly in the opposite direction of the miRNA (miRhub p-value<0.05; **Figure 4B**). To account for the genetic cofounders that emerge when comparing primary tumors of one patient to non-tumor tissue from another patient, we also performed a differential miRNA expression analysis between matched tissues (n=8). Of the 17 miRNAs identified above, 15 are still significantly altered when the analysis is restricted to matched samples (**Figure 4C**).

**Figure 4:**
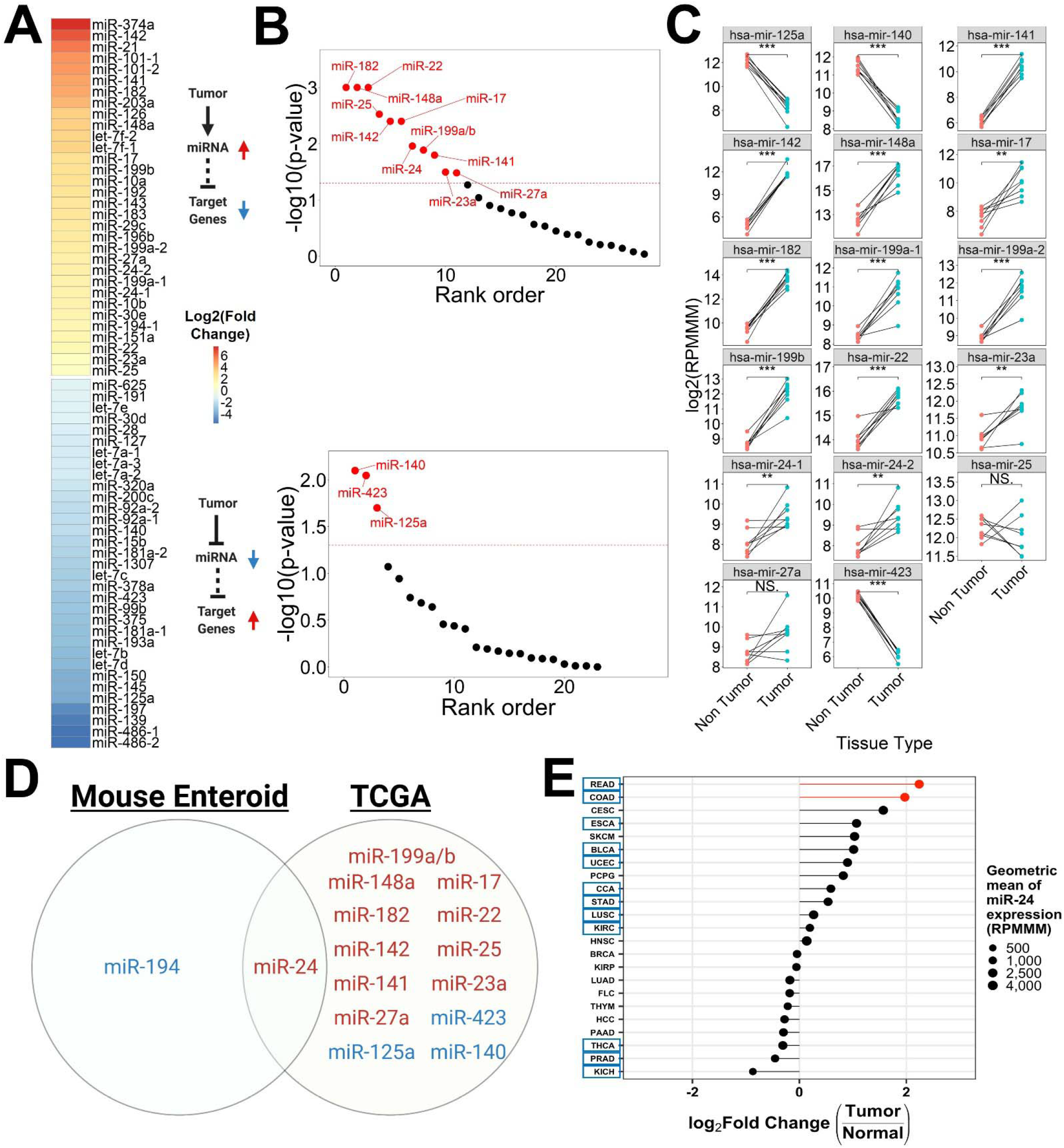
miR-24-3p is a candidate regulator of gene expression in human CRC. (**A**) Heatmap showing the log2 fold change for miRNAs differentially expressed (>1000 RPMMM in either condition, fold change >1.5x, p-adj<0.05) between TCGA primary colon adenocarcinoma (n=371) and non-tumor tissue (n=8). Color intensity represents the log2 fold change. (**B**) Plot of the -log10 (p-value) of target site enrichment, calculated by miRhub (cons2) for each differentially expressed miRNA from (**A**), using the list of genes that are differentially expressed (DESeq2 expression >1000 normalized counts in either condition, fold change >1.5x, p-adj<0.05) in the opposite direction of the miRNA. MiRNAs within the same family were grouped together under the same name. MiRNAs with target site enrichment p-value<0.05 shown in red. (**C**) Expression (log2 RPMMM) of the 17 miRNAs from (**B**) in matched TCGA primary colon adenocarcinoma (n=8) and non-tumor (n=8) tissue (two-tailed Welch t-test). Lines connect tissue samples collected from the same patient. (**D**) Venn diagram for miRNAs of interest identified by the mouse enteroid and TCGA analyses (Created with BioRender.com). MiRNAs in red are upregulated. MiRNAs in blue are downregulated. Paralogs are listed as one miRNA. (**E**) Log2 fold change of miR-24-3p expression (RPMMM) across TCGA tumor types (n=23). Colon (COAD) and rectal (READ) adenocarcinomas in red. Circle size represents the geometric mean (RPMMM) of miR-24-3p for each tumor type. Tumor types highlighted by blue boxes have Benjamini-Hochberg padj<0.05. *p<0.05, **p<0.01, ***p<0.001.

Of these 15 miRNAs that are candidate key regulators of gene expression in human CRC, only miR-24-3p was also identified as a candidate regulator in the mouse enteroid analyses (**Figure 4D**). We divided TCGA primary colon tumors into genotype bins that corresponded to the mutational combinations generated in our enteroid models. For genotypes with sample size was >3 tumors, we observe a significant elevation in miR-24-3p expression compared to non-tumor controls (**Table S3**). For genotypes with sample sizes less than 4, we are limited in our ability to confidently identify differentially expressed miRNAs given the high cellular and genetic heterogeneity of the tissues. We next assessed changes in miR-24-3p expression across 23 different tumor types relative to their corresponding non-tumor tissue. We find that 12/23 tumor types have a significant alteration in miR-24-3p expression (**Figure 4E**). Of these, rectal adenocarcinoma (READ) and colon adenocarcinoma (COAD) have the highest upregulation of miR-24-3p (**Figure 4E**), indicating that miR-24-3p upregulation is strongest in CRC.

### Reduction of miR-24-3p increases apoptosis in HCT116 cells

We hypothesized that miR-24-3p promotes colon tumor phenotypes. To evaluate this hypothesis, we performed loss-of-function studies in HCT116 cells, which is derived from a microsatellite instable human colon tumor with mutations in *CTNNB1, KRAS*, and *TGFBR3*. Specifically, we treated HCT116 cells with a miR-24-3p locked nucleic acid (LNA) inhibitor, which led to significantly reduced detection of miR-24-3p (**Figure 5A**). HCT116 cultures treated with a miR-24-3p inhibitor exhibit a significant reduction in cell number compared to mock and scramble controls (**Figure 5B**). We also observe a significant reduction in the number of metabolically active, viable cells as determined by the CellTiter-Glo assay (**Figure 5C**). CellTiter-Glo experiments were repeated in three additional cell lines with various degrees of effect on cell viability (**Supplementary Figure 4**). Subsequent studies continued to utilize HCT116 cells as we observed the strongest effect on cell viability in this cell context.

**Figure 5:**
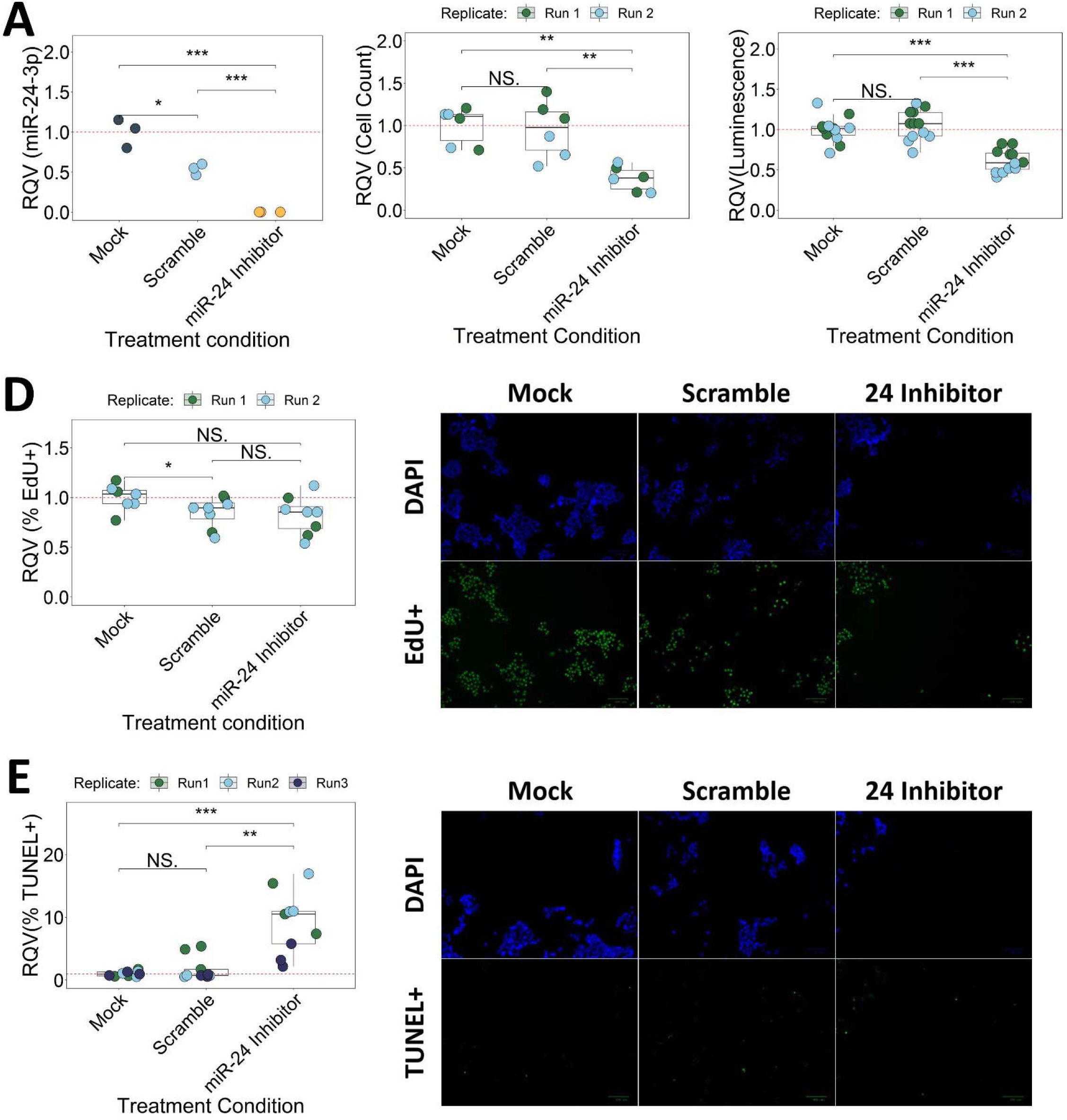
Inhibition of miR-24-3p increases apoptosis in HCT116 cells. (**A**) MiR-24-3p expression from RT-qPCR following mock, 100 nM scramble, or 100 nM miR-24 inhibitor treatment of HCT116 cells. Significance determined by two-tailed Student’s t-test. Cell count (**B**), CellTiter-glo (**C**), EdU incorporation (**D**), and TUNEL (**E**) assays following mock, 100 nM scramble, or 100 nM miR-24 inhibitor treatment in HCT116 cells. Signficance determined by two-sided Wilcoxon test. Results reported relative to average mock control. Color of data points represents experimental replicate. *p<0.05, **p<0.01, ***p<0.001.

Next we asked whether the change in cell viability was caused by differences in the rate of cell proliferation, cell death, or both. To evaluate changes in proliferation, we performed an EdU incorporation assay. Our analysis shows a significant decrease in the number of DAPI+ cells (**Supplementary Figure 5A**) but does not identify a significant change in the percentage of EdU+ cells after treatment with miR-24-3p inhibitor relative to mock or scramble controls (**Figure 5D**). To evaluate changes in apoptosis, we performed a TUNEL assay. HCT116 cells treated with a miR-24-3p inhibitor display a significant decrease in the number of DAPI+ cells (**Supplementary Figure 5B**) and an increase in the percentage of TUNEL+ cells compared to mock and scramble controls (**Figure 5E**). Thus, we conclude that miR-24-3p promotes CRC cell viability at least in part through suppression of apoptosis (and not through increased proliferation).

### miR-24-3p inhibition decreases mouse enteroid survival

To further validate the role of miR-24-3p in regulating cell survival in the intestine, we next examined the effects of miR-24-3p inhibition on the growth and viability of mouse enteroids. Jejunal crypts were isolated from WT B62J mice and cultured ex vivo to establish enteroids, which were treated with either a miR-24-3p LNA inhibitor or scramble control for a total of five days. Enteroid cultures treated with the miR-24-3p inhibitor exhibit significant (∼33%) reduction in the number of enteroids (**Figure 6A, Supplementary Figure 6A**). However, enteroids treated with a miR-24-3p inhibitor do not exhibit a significant difference in enteroid size relative to those treated with the scramble control (**Figure 6B, Supplementary Figure 6B**). Taken together, these results provide further support that miR-24-3p promotes cell survival of intestinal epithelial cells.

**Figure 6:**
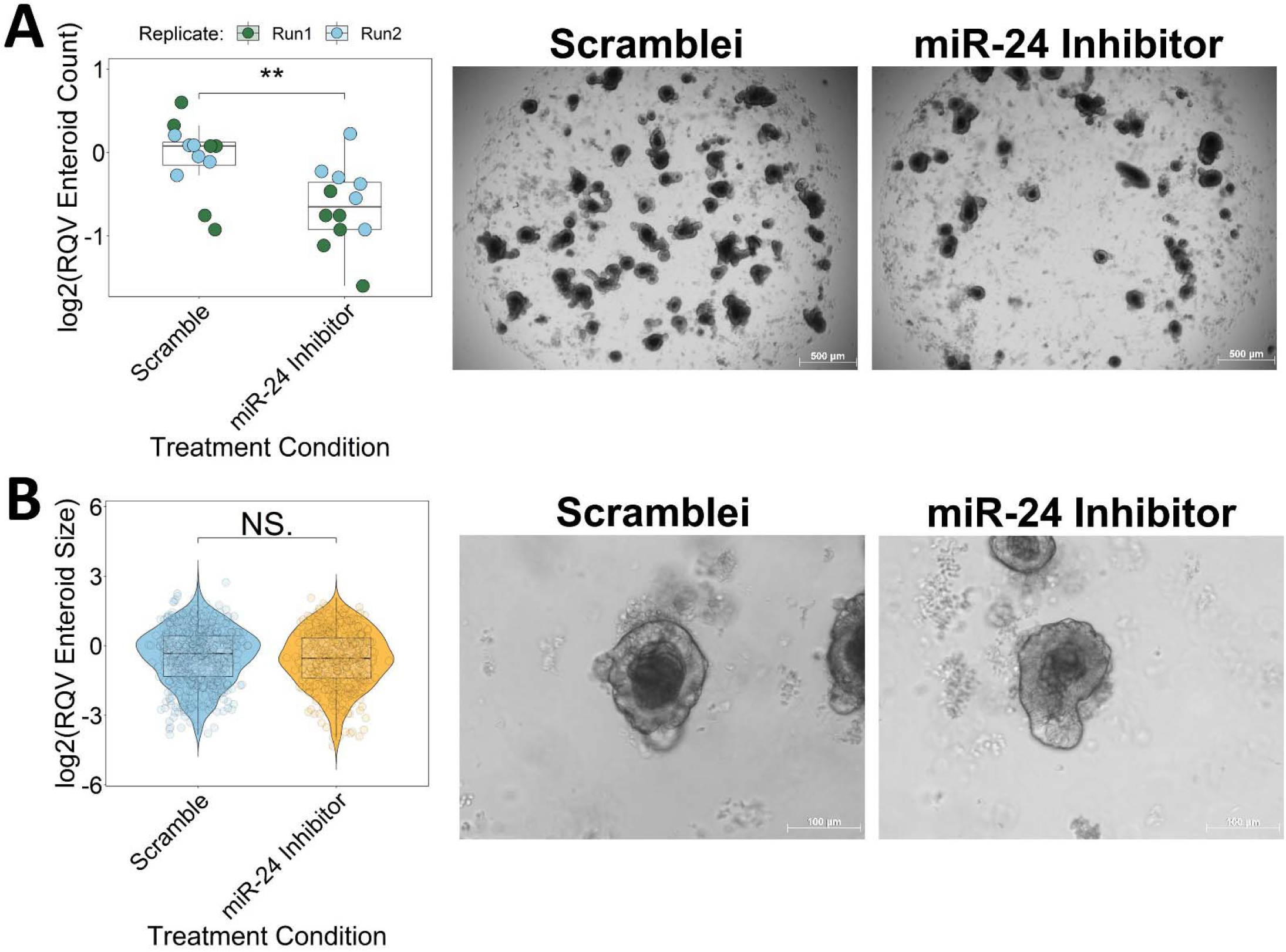
miR-24-3p inhibition decreases mouse enteroid survival. (**A**) Number of WT enteroids following scramble or miR-24 inhibitor treatment. Significance determined by two-tailed Welch t-test. Data reported relative to scramble average. Color of data points represents experimental replicate. (**B**) Violin plot of enteroid size across experimental replicates following scramble or miR-24 inhibitor treatment. Significance determined by two-tailed Welch t-test. Data reported relative to average scramble control. *p<0.05, **p<0.01, ***p<0.001.

### HMOX1 and PRSS8 are post-transcriptionally regulated by miR-24 in CRC

To identify candidate gene targets by which miR-24-3p exerts its function in CRC, we treated HCT116 cells with miR-24-3p LNA inhibitor or scramble control. After 48 hrs, we isolated RNA from these cells and performed an RNA-seq analysis to identify genes that change in response to miR-24-3p inhibition (**Table S4**). We reasoned that direct target genes should be inversely correlated with miR-24-3p; therefore, we focused our subsequent analyses on the 222 genes that are significantly elevated (expression above 500 normalized counts in either condition, p-adj<0.05, Fold change > 0) in response to miR-24-3p inhibition (**Figure 7A**). Of these genes, 70 are predicted miR-24-3p targets (**Figure 7B**). We performed a KEGG pathway analysis using Enrichr^52-54^ (**Figure 7C**), which reveals that up-regulated genes with predicted miR-24-3p targets are enriched in apoptosis and ferroptosis pathways (two different forms of cell death). Notably, nine of the 70 predicted miR-24-3p target genes are also significantly downregulated (DESeq2; average expression > 1000 normalized counts, >1.5x fold change, p-adj<0.05) in TCGA colon tumor relative to non-tumor tissue (**Figure 7B**), including *HMOX1* and *PRSS8*, which exhibit the highest upregulation among the nine (**Figure 7D**). Moving forward, we focused on *HMOX1* and *PRSS8*, which have been shown previously to regulate cell survival in various cancer contexts^55-58^.

**Figure 7:**
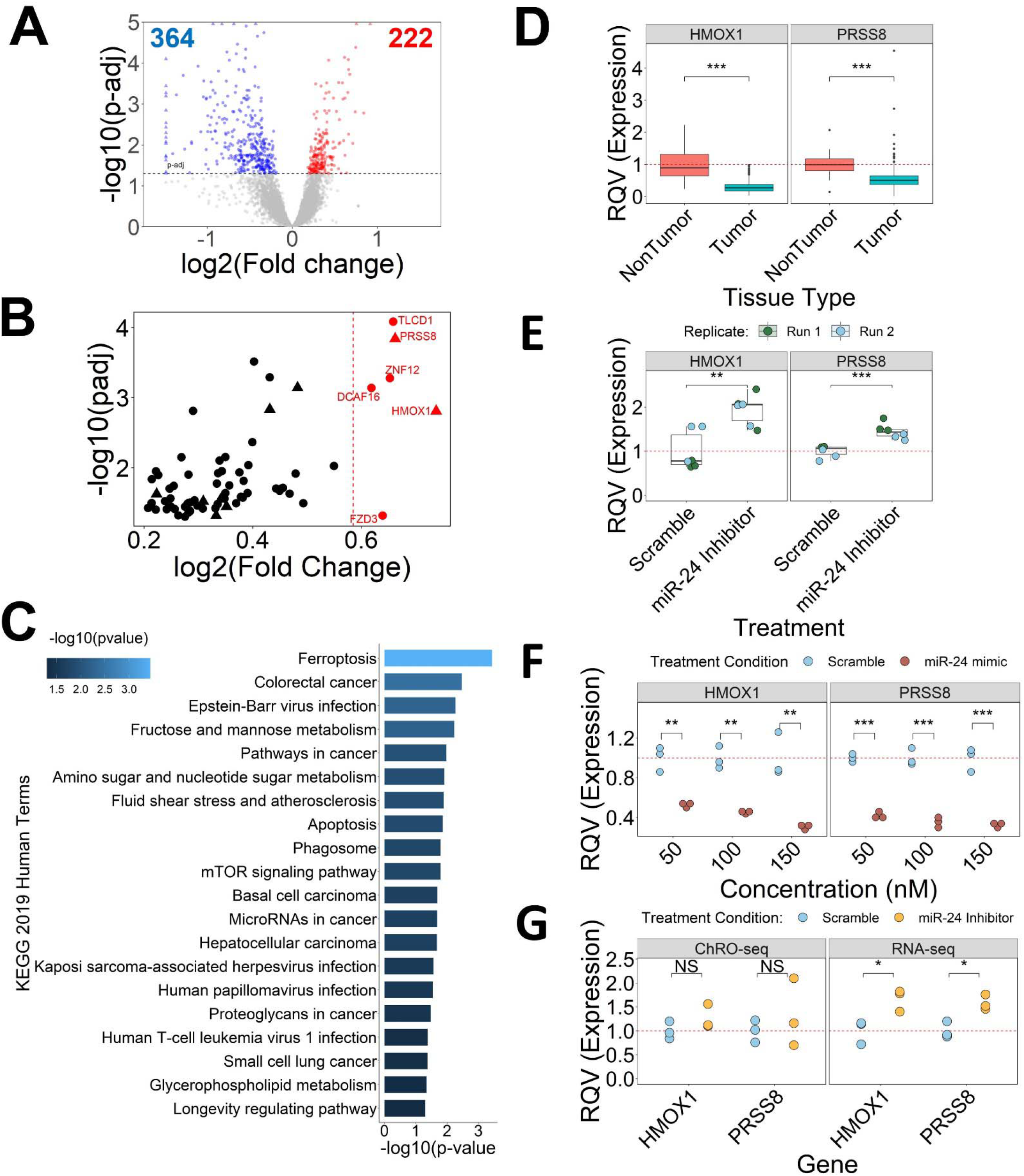
*HMOX1* and *PRSS8* are post-transcriptionally regulated by miR-24-3p. (**A**) Volcano plot showing differentially expressed genes in HCT116 treated with a 100 nM miR-24 inhibitor relative to scramble control. Genes filtered for expression >500 normalized counts in either condition. Horizontal dashed line represents p-adj cutoff of 0.05 (DESeq2). (**B**) Scatterplot of predicted miR-24-3p target genes that are upregulated (DESeq2 p-adj<0.05, >500 normalized counts in either condition) following miR-24-3p inhibition (n=70). Vertical red line represents 1.5x fold change. Genes in red exhibit >1.5x fold change (n=6). Genes also significantly downregulated in TCGA tumor tissue compared to non-tumor are represented by triangles. Remaining genes represented by circles. (**C**) KEGG pathway enrichment analysis of 70 genes in (**B**). Pathways with p-value <0.05 represented in figure. Color represents the -log10 p-value. (**D**) Normalized expression from small RNA-seq of *HMOX1* and *PRSS8* in TCGA colon tumor relative to non-tumor tissue. Significance determined by two-sided Wilcoxon test. (**E**) RT-qPCR for *HMOX1* and *PRSS8* following 100 nM miR-24 inhibitor or scramble treatment in HCT116 cells. Significance determined by two-tailed Welch t-test. Color of data points represents experimental replicate. (**F**) RT-qPCR for *HMOX1* and *PRSS8* following HCT116 treatment with 50, 100 or 150 nM miR-24 mimic or scramble. Significance determined by two-tailed Student’s t-test. A non-parametric test was applied (two-sided Wilcoxon test), but significance couldn’t be achieved due to low sample size. (**G**) DESeq2 normalized RNA-seq and ChRO-seq counts for *HMOX1* and *PRSS8* expression following HCT116 treatment with scramble or miR-24 inhibitor. Significance determined by two-tailed Welch t-test. *p<0.05, **p<0.01, ***p<0.001.

As an independent validation of the RNA-seq analysis, we performed RT-qPCR analysis for *HMOX1* and *PRSS8* using RNA from HCT116 cells treated with a miR-24-3p LNA inhibitor or scramble control. As expected, both genes exhibit a significant elevation in miR-24-3p inhibitor treated cells compared to control (**Figure 7E**). We also treated HCT116 cells with 50, 100, or 150 nM of miR-24-3p mimic or scramble control. Consistent with expectation, we observe a dose-dependent decrease in *HMOX1* and *PRSS8* expression (**Figure 7F**). These results support the model that miR-24-3p regulates *HMOX1* and *PRSS8* in CRC cells.

Regulation of *HMOX1* and *PRSS8* by miR-24-3p could occur through the canonical miRNA post-transcriptional gene targeting or by indirectly controlling the transcriptional activity of the two genes. To distinguish between these two possibilities, we leverage length extension chromatin run-on sequencing (leChRO-seq)^28^ to assess changes in *HMOX1* and *PRSS8* transcription following miR-24-3p inhibition. Solely post-transcriptionally regulated genes will exhibit similar transcription (dectected by leChRO-seq) between scramble and miR-24-3p inhibitor treated cells, but altered steady-state gene expression (measured by RNA-seq). We show that *HMOX1* and *PRSS8* are transcribed at a similar rate between miR-24-3p inhibitor and scramble treated HCT116 cells, as determined by leChRO-seq (**Figure 7G, Table S5**). However, both genes do exhibit a significant elevation at the mRNA level as detected by RNA-seq (Figure 7G). Together, this data supports that miR-24-3p post-transcriptionally regulates *HMOX1* and *PRSS8* in a CRC context.

## Discussion

In this study we leveraged genetically modified mouse enteroids to characterize the impact of different combinations of CRC driver mutations on miRNA expression. We show that each of the genotypes investigated result in distinct miRNA profiles, with the exception of *Rspo3* (R) mutant enteroids which are comparable to wild-type (WT). The latter finding is consistent with previous studies that have shown R mutant enteroids exhibit similar RNA-seq profiles and cell morphology to WT enteroids^31^. We also define separate modules of miRNAs, each of which exhibits a unique pattern of expression across genotypes. We establish a publicly accessible resource, called ME-MIRAGE (https://jwvillan.shinyapps.io/ME-MIRAGE/), that allows users to evaluate mutation-specific relationships between miRNAs and genes. This database is a novel resource that provides information regarding the miRNA-mediated mechanisms by which combinations of somatic mutations can drive inter-tumor heterogeneity in CRC.

We highlight miR-375-3p as a mutation-dependent miRNA by showing that its expression is most strongly affected in the Smad4^KO^ context. Interestingly, *Apc* (A) mutant enteroids exhibit significantly reduced (DESeq2 fold change > 1.5x, padj < 0.05) miR-375-3p expression compared to WT, however the magnitude of the decrease is much smaller relative to KRS and KRPS enteroids. This suggests that inhibition of miR-375-3p in CRC is strongly, but not solely, driven by changes in Tgf-B signaling. Studies in other cell contexts suggest that miR-375-3p can regulate Tgf-B signaling^59,60^. Future directions may explore the downstream effects and potential feedback mechanisms by which miR-375-3p can regulate Tgf-B signaling. Additionally, previous studies in colon, stomach, and liver cancers have established miR-375-3p as a tumor suppressor^47,61,62^. This is in line with our recent report that shows miR-375 inhibits cell proliferation and migration in fibrolamellar carcinoma^46^ and suppresses proliferation in intestinal stem cells^63^. In CRC, we suggest that a miR-375-3p mimic could be a candidate therapeutic approach especially for patients with somatic mutations that inhibit Tgf-B signaling.

The limited literature on the functional role of miR-24-3p in CRC offers mixed conclusions on whether miR-24-3p is an upregulated oncogenic miRNA^64,65^ or a downregulated tumor suppressor^66,67^. This is likely due to a combination of pleiotropy in miR-24-3p function and the differences in experimental approaches across studies. In this study, we leverage multiple cell models in addition to TCGA data to support that miR-24-3p is upregulated in CRC and can function as an oncogenic miRNA. MiR-24-3p is located on the same pri-miRNA transcript as miR-27a-3p and miR-23a-3p^68^. We found that all three miRNAs are significantly elevated in TCGA primary colon tumor tissue and that genes downregulated in CRC are enriched for predicted targets of each of the miRNAs. MiR-27a-3p and miR-23a-3p also exhibit elevated expression (albeit not always statistically significant) in multiple mutant enteroids relative to WT. This upregulation in the miR-23a/miR-24/miR-27a cluster across datasets supports the literature that miR-24-3p is elevated in CRC. Finally, functional studies in HCT116 cells and enteroids demonstrate that inhibition of miR-24-3p suppresses CRC tumor cell apoptosis. Pleiotropy may explain the varied responses we observed in other CRC cell lines to miR-24-3p inhibition (**Supplementary Figure 4**). These results would suggest that, although miR-24-3p is upregulated in a wide range of genetic contexts, there is heterogeneity in the responsiveness of CRC cells to miR-24-3p inhibition.

We identify multiple genes that are up-regulated due to loss of post-transcriptional suppression after inhibition of miR-24-3p, most notably *HMOX1* and *PRSS8*, which are also prominently downregulated in TCGA colon tumors. Given miR-24-3p inhibition leads to increased apoptosis, we predict that HMOX1 and PRSS8 function as tumor suppressors in CRC. While PRSS8 has clearly been shown to promote apoptosis in multiple cancer contexts^57,58^, HMOX1 in regulating apoptosis appears to vary across tissues^55,69,70^. While the role of HMOX1 in the colon remains to be thoroughly evaluated, studies show that HMOX1 can function as a tumor suppressor in CRC by inhibiting tumor invasion^71^ and metastasis^72^. Here we suggest that HMOX1 may also function as a tumor suppressor by increasing apoptosis, which merits more detailed future investigation.

In our KEGG 2019 pathway enrichment analysis of upregulated genes after miR-24-3p inhibitor treatment we identified ferroptosis, a form of cell death induced by excessive iron-induced lipid peroxidation. Cell count and CellTiter-glo analyses of HCT116 cells treated with a miR-24-3p inhibitor and ferroptosis inhibitor, ferrostatin, reveals no partial recovery of cell number at increasing concentrations of ferrostatin (**Supplementary Figure7**). One potential explanation for this data is that the poor stability of ferrostatin^73^ prevented effective inhibition of ferroptosis. Another possibility is that miR-24-3p inhibition primed cells to undergo ferroptosis, but without the proper induction of ferroptosis, we do not observe a change in cell number at higher concentrations of ferrostatin. Treating cells with cisplatin, or another platinum-based therapy like oxaliplatin which is commonly used to treat CRC patients, may be an appropriate stimulus as cisplatin has been shown to induce apoptosis and ferroptosis in HCT116 cells^74^. If miR-24-3p does inhibit ferroptotic cell death, then treating CRC patients with a miR-24-3p inhibitor in conjunction with oxaliplatin may increase the efficacy of treatment and increase patient survival. We believe this idea merits further investigation.

Our results provide insight into the mechanisms by which somatic mutations alter miRNA profiles and how this can contribute to inter-tumor heterogeneity. Future studies in the field can build on our work by incorporating somatic mutations in genes that stratify established CRC subtypes^3^ and observing changes in miRNA profiles. Further understanding of mutation-specific alterations to oncogenic and tumor suppressive miRNAs will be important for determining which miRNA-based therapeutics are most effective in different mutational contexts. Additionally, we hope to characterize how combinations of somatic mutations affect pri-miRNA transcription to elucidate the transcriptional programs that contribute to changes in mature miRNA profiles. Ultimately, the identification of mutation-specific miRNAs will be important for identifying candidate miRNA therapeutics and the overall advancement of precision medicine for CRC patients.

## Experimental Procedures

### Generation of genetically modified mouse enteroids

The proximal half of the small intestine was isolated from 5-6 week-old C57BL/6 mice for crypt isolation. Cells were plated in Matrigel and grown for 3-4 weeks. Enteroids were then dissociated and transfected using the necessary Cre/CRISPR gene editors.

#### A, B, AKP, BKP

Cells with *Apc*^Q883*^ mutation (A) and *Ctnnb1*^S33F^ (B) were generated using CRISPR base editing as described in Schatoff et al. (2019)^29^. Selection for *Apc*^Q883*^ and *Ctnnb1*^S33F^ mutants was performed by culturing cells in the absence of RSPO1. *Kras*^G12D^ (K) and *Trp53*^-/-^ (P) mutations were generated using enteroids from the conditional LSL-*Kras*/*p53*fl/fl mouse model, as described in Dow et al., (2015)^30^. Cre was introduced to enteroids by transfection. Cells with *Trp53*^-/-^ mutation were selected for by treating cells with 10 µM Nutlin3. To ensure *Kras*^G12D^ mutation, cells were then cultured in the absence of EGF.

#### KRP, KRS, KRPS

*Kras*^G12D^ mutations (K) were generated by using enteroids derived from the *Kras*^LSL-G12D^ conditional model as described in Jackson et al. (2001)^75^. Cre was introduced to enteroids by transfection. Cells with the Kras^LSL-G12D^ allele were selected for by adding 1µM gefitinib to the culturing media. Cells with *Ptprk*-*Rspo3* fusion (R) were generated via CRISPR/Cas9 chromosome rearrangement as described in Han et al., (2017)^31^. Selection for *Ptprk*-*Rspo3* mutants was done by culturing cells in the absence of RSPO1. *Trp53*^-/-^ (P)and *Smad4*^KO^ (S) mutations were generated using CRISPR/Cas9 and single guide RNAs (sgRNA) as described in Han et al., (2020)^12^. Selection for *Trp53*^-/-^ cells was completed by adding 5 µmol/L Nutlin-3 to the culturing media. Selection for *Smad4*^KO^ cells was completed by adding 5 ng/mL TGFB1 to the culturing media.

### Trizol LS RNA isolation

Cells were treated with 250 µL of cold 1X NUN Lysis Buffer (20 mM HEPES, 7.5 mM MgCl2, 0.2 mM EDTA, 0.3 M NaCl, 1 M Urea, 1% NP-40, 1mM DTT, and 50 units/mL SUPERase In RNase Inhibitor (ThermoFisher Scientific, Waltham, MA), and 1X 50X Protease Inhibitor Cocktail (Roche, Branchburg, NJ)). Lysate was vortexed vigorously for 1 minute to physically lyse cells. Samples were incubated for 30 minutes in Thermomixer C at 12°C at 1500 rpm. Chromatin was pelleted out by centrifuging samples for 12,500 xg for 30 minutes at 4°C. Supernatant containing RNA was removed from the tube and added to clean 1.5 mL centrifuge tube along with 750 µL Trizol LS (Life Technologies, 10296–010). Samples were vortexed and stored at -80°C until RNA isolation. Samples were thawed and allowed to incubate for 5 minutes. 200 µL of chloroform was added to each tube and vortexed for 20 seconds. Following a three-minute incubation, samples were centrifuged at 17,000 xg, 4°C for 5 minutes. Aqueous layer was transferred to clean 1.5 mL centrifuge tube containing 2.5 µL of GlycoBlue. 1 mL of ice cold, 100% ethanol was added to aqueous phase and samples were then vortexed. Samples were then centrifuged at 17,000 xg at 4°C for 15 minutes. Supernatant was removed and pellet was washed with 75% ice cold ethanol. Samples were vortexed and RNA was pelleted by centrifuging at 17,000 xg at 4°C for 5 minutes. Supernatant was removed and RNA pellets were allowed to dry for 10 minutes at room temperature. RNA was resuspended in 30 µL of RNase-free water.

### Small RNA library preparation and sequencing

Total RNA was isolated using the Total RNA Purification Kit (Norgen Biotek, Thorold, ON, Canada) according to manufacturer’s instructions or Trizol LS method described above. RNA purity and concentration was determined using the Nanodrop 2000 (Thermo Fisher Scientific, Waltham, MA). RNA integrity was quantified using the 4200 Tapestation (Agilent Technologies, Santa Clara, CA) or Fragment Analyzer Automated CE System (Advanced Analytical Technologies, Ankeny, IA). Libraries were prepared at the Genome Sequencing Facility of the Greehey Children’s Cancer Research Institute (University of Texas Health Science Center, San Antonio, TX) using the CleanTag Small RNA Library Prep kit (TriLink Biotechnologies, San Diego, CA). Libraries were then sequenced on the HiSeq2000 platform (Illumina, San Diego, CA).

### RNA library preparation and sequencing

Total RNA was isolated using the Total RNA Purification Kit (Norgen Biotek, Thorold, ON, Canada) according to the manufacturer’s instructions or using the Trizol LS method described above. RNA purity and concentration was determined using the Nanodrop 2000 (Thermo Fisher Scientific, Waltham, MA). RNA integrity was quantified using the 4200 Tapestation (Agilent Technologies, Santa Clara, CA) or Fragment Analyzer Automated CE System (Advanced Analytical Technologies, Ankeny, IA). Libraries were prepared using the NEBNext Ultra II Directional Library Prep Kit following Ribosomal Depletion (mouse enteroid RNA) or PolyA enrichment (HCT116 RNA) at the Cornell Transcriptional Regulation and Expression Facility (Cornell University, Ithaca, NY). Libraries were then sequenced using the NextSeq500 platform (Illumina, San Diego, CA).

### Small RNA-seq Analysis

Read quality was assessed using FastQC. Trimming, mapping and quantification was performed using miRquant 2.0 as described in Kanke et al., (2016)^32^. In short, reads were trimmed using Cutadapt, aligned to the genome using Bowtie and SHRiMP, and aligned reads were quantified and normalized using DESeq2^50^. We accounted for sequencing batch, RIN, and genotype in our model. <u>Defining groups of miRNAs with similar patterns of expression across genotypes:</u> Raw miRNA count matrices produced by miRquant were analyzed using a likelihood ratio-test from DESeq2. miRNA annotations in the 5000s are degradation products and removed from the analysis, and miRNAs with an adjusted p-value greater than 0.05 and baseMean expression less than 500 were discarded. An rlog transformation was applied to the raw counts and batch effects were removed using the limma function removeBatchEffects. Clusters of miRNAs with similar expression patterns were identified using the DEGreport (v1.26.0) function degPatterns (minc = 5). Only clusters containing greater than five miRNAs were considered. <u>Fold change heatmaps:</u> Transformation and batch correction of miRNA expression and grouping of miRNAs is described above. Normalized expression of miRNAs for each mutant enteroid sample was subtracted from average WT expression and heatmaps were made using the R package pheatmap (v1.0.12).

### RNA-seq Analysis

Read quality was assessed using FastQC. RNA-seq reads were aligned to either the mm10 genome release for mouse enteroids or the hg38 genome release for the human HCT116 cells using STAR (v2.7.9a). Quantification was performed with Salmon (v1.4.0) using the GENCODE release 25 annotations. Normalization and differential expression analyses were performed utilizing DESeq2 (v1.30.1). We accounted for cell culture batch effects, RIN, and genotype in our model. Enrichr was used for KEGG pathway analysis as described in Chen et al. (2013)^52^. <u>miRhub analysis</u> was performed as described in Baran-Gale et al. (2013)^51^. In short, miRhub scans input gene lists for miRNA binding sites defined by TargetScan v5.2^76^. For our analyses, we filtered for binding sites that are conserved in mice and at least one of the following species (cons1): human, rat, dog and/or chicken. miRNA-gene scores were generated based on seed sequence strength, conservation, and frequency of target sites in the 3’-UTR while controlling for 3’ UTR length. These scores were added together for each miRNA to generate a cumulative value that represents the miRNA targeting score. A Monte Carlo simulation repeated this analysis 1000x using lists of randomly selected genes. An empirical p-value was then calculated for each miRNA by comparing the targeting score from input gene lists to the targeting scores calculated calculated using the lists consisting of randomly selected genes.

### Quantitative PCR

Total RNA from HCT116 cells was extracted using the Total RNA Purification Kit (Norgen Biotek, Thorold, ON, Canada) according to manufacturer’s instructions. Reverse-transcription for <u>miRNA expression</u> was performed using the Taqman MicroRNA Reverse Transcription Kit (ThermoFisher Scientific, Waltham, MA). Quantification of miRNA expression was done using the TaqMan Universal PCR Master Mix (ThermoFisher Scientific, Waltham, MA). miRNA expression was normalized to U6 (assay ID: 001973). miRNA Taqman assays: miR-375-3p (assay ID: 000564), miR-24-3p (assay ID: 000402). Reverse-transcription for <u>gene expression</u> was performed using the High-Capacity RNA-to-cDNA kit (ThermoFisher Scientific, Waltham, MA). Quantification of gene expression was done using the TaqMan Gene Expression Master Mix (ThermoFisher Scientific, Waltham, MA). Gene expression was normalized to *RPS9* (assay ID: Hs02339424_g1). Gene Taqman assays: *HMOX1* (assay ID: Hs01110250_m1), *PRSS8* (assay ID: Hs00173606_m1), *Rps9* (assay ID: Mm00850060_s1), *Fn1* (assay ID: Mm01256744_m1), *Col1a1* (assay ID: Mm00801666_g1). Measurements were taken using the BioRad CFX96 Touch Real Time PCR Detection System (Bio-Rad Laboratories, Richmond, CA).

### The Cancer Genome Atlas (TCGA) Analysis

#### Data Download

RNA-seq High Throughput Sequencing (HTSeq) counts files for 382 primary colon tumor and 39 solid normal tissue samples was downloaded using the NIHGDC Data Transfer Tool. Normalization and differential expression were identified using DESeq2. For our miRhub analysis, we filtered for binding sites that are conserved in humans and at least two of the following species (cons2): mouse, rat, dog and/or chicken. miRNA quantification files, that used mirbase21, for 371 primary colon tumor and 8 solid normal tissue samples were also downloaded using NIHGDC Data Transfer Tool. Of the 371 colon tumor samples with miRNA data, 326 had simple somatic mutation (TCGA v32.0) and copy number variation (CNV; TCGA v31.0) information. Tumor samples were assigned *APC* (A), *TP53* (P), and *SMAD4* (S) mutations if they contained a non-synonymous mutation and/or CNV loss for a given gene. For A, P, and S designations, samples with a CNV gain and a non-synonymous mutation were not included. Mutations in *CTNNB1* (B) and *KRAS* (K) were assigned to tumor samples with a non-synonymous mutation and/or CNV gain for a given gene. For B and K designations, samples with a CNV loss and a non-synonymous mutation were not included.

#### TCGA small RNA-seq across cancer types

Small RNA sequencing expression data was downloaded from TCGA for 23 tumor types using the R package TCGA-assembler. Expression was reported as the reads per million mapped to miRNAs (RPMMM). Log2 fold change was calculated by dividing the tumor expression by the expression in non-tumor tissue followed by log2 transformation.

### Mouse enteroid culture

Crypts from the jejunum of 3-5 month old male B62J mice were isolated as described in Peck et al., (2017)^63^. Isolated crypts were plated in Reduced Growth Factor Matrigel (Corning, Corning, NY, catalog #: 356231) on Day 0. Advanced DMEM/F12 (Gibco, Gaithersburg, MD, catalog #: 12634-028) was used for culture and supplemented with GlutaMAX (Gibco, Gaithersburg, MD, catalog #:35050-061), Pen/Strep (Gibco, Gaithersburg, MD, catalog #:15140), HEPES (Gibco, Gaithersburg, MD, catalog #:15630-080), N2 supplement (Gibco, Gaithersburg, MD, catalog #:17502-048), 50 ng/mL EGF (R&D Systems, Minneapolis, MN, catalog #: 2028-EG), 100 ug/mL Noggin (PeproTech, Rocky Hill, NJ, catalog #: 250-38), 250 ng/uL murine R-spondin (R&D Systems, catalog #: 3474-RS-050), and 10 mM Y27632 (Enzo Life Sciences, Farmingdale, NY, catalog #:ALX270-333-M025) <u>miR-24-3p LNA inhibitor treatment:</u> Cells were transfected with hsa-miR-24-3p miRCURY LNA miRNA Power Inhibitor (Qiagen, Germantown, MD, catalog #: YI04101706-DDA) or Power Negative Control A (Qiagen, Germantown, MD, catalog #: YI00199006-DDA) to a final concentration of 500 nM on Day 0 using gymnosis. Media was changed and cells were treated with 250 nM miR-24 LNA inhibitor or scramble. Cells were harvested and fixed in 4% (v/v) paraformaldehyde on Day 5. <u>Tgf-B treatment:</u> Recombinant Human TGF-B1 (PeproTech catalog #: 100-21) was added to enteroid media on Day 0 for final concentration of 0, 0.5, or 1 ng/mL. Enteroids were harvested on Day 3.

### Cell Line Transfection

All cell lines were plated in DMEM+10% FBS media. HCT116 cells were plated at a density of 3,400 cells/well in a 96-well plate. Caco-2 cells were plated at a density of 20,000 cells/well in a 96-well plate. HT-29 cells were plated at a density of 3,400 to 6,800 cells/well in a 96-well plate. SW48 cells were plated at a density of 10,000 cells/well in a 96-well plate. Cells incubated for 24 hours in a 37 °C incubator. Cells were transfected with hsa-miR-24-3p miRCURY LNA miRNA Power Inhibitor (Qiagen, Germantown, MD, catalog #: YI04101706-DDA) or scramble control to a final concentration of 100 nM using Lipofectamine 3000 (ThermoFisher Scientific, Waltham, MA, catalog #: L3000-008) according to manufacturer’s instructions. Either Power Negative Control A (Qiagen, Germantown, MD, catalog #: YI00199006-DDA) or Negative Control miRCURY LNA miRNA Mimic (Qiagen, Germantown, MD, catalog #: YM00479902-AGA) was used for scramble control. After 24-hours, media was replaced with complete media. <u>Ferrostatin-1 treatment:</u> At the time of LNA transfection, cells were also treated with 0, 0.5, 2, 5, or 10 µM Ferrostatin-1 (Sigma-Aldrich, St. Louis, MO, catalog #: SML0583-5MG). After 24-hours, media was replaced with complete media. Cells were harvested 48-hours post-transfection.

### Cell count assay

48-hours following transfection, cells in 96-well plate were washed with PBS and treated with 50 µL trypsin. Cells incubated for 5 minutes in 37 °C incubator. Cells were resuspended using 150 µL complete media and transferred to clean 1.5 mL Eppendorf tubes. Cell concentration was calculated by adding 10 µL of cell suspension to chip for Biorad TC20 Automated Cell Counter (Bio-Rad Laboratories, Richmond, CA).

### CellTiter-Glo assays

48-hours following transfection, cells in 96-well plate were incubated at room temperature for 30 minutes. 100 µL of room temperature CellTiter-Glo reagent (Promega, Madison, WI) was added to each well and placed on cell rocker for 2 minutes to lyse the cells. Afterwards, plate was incubated at room temperature for 10 minutes. Luminescent signal was quantified using a Synergy 2 Microplate Reader (Biotek, Winooski, VT; area scan; Integration = 0:00:50; Sensitivity = 135).

### EdU Assay

48-hours following transfection, cells in 96-well plate were incubated with 10 µM EdU at 37 °C in complete media for 1 hour. Cells were then fixed with 4% paraformaldehyde for 20 minutes at room temperature and permeabilized using 0.5% Triton X-100 in PBS for 20 minutes. The Invitrogen Click-iT Plus EdU AlexaFluor 488 Imaging Kit (Invitrogen, Waltham, MA, C10637) was used to detect EdU according to manufacturer’s instructions. Nuclei were stained using DAPI (ThermoFisher Scientific, Waltham, MA, catalog #: D1306) and imaged using ZOE Fluorescent Cell Image (Bio-Rad Laboratories, Richmond, CA). Images were analyzed using FIJI. For EdU positive cells, threshold value was set to 10. For analyzing particles, counted those particles with size = 250-Infinity and circularity = 0.4-1.

### TUNEL Assay

48-hours following transfection, cells in 96-well plate were washed twice with PBS and fixed using 4% paraformaldehyde for 15 minutes at room temperature. Permeabilization was performed by using 0.5% Triton X-100 in PBS for 20 minutes. Cells were washed twice with deionized water. Positive control wells were treated with 1X DNase I, Amplification Grade (ThermoFisher Scientific, Waltham, MA, catalog #: 18068-015) solution according to manufacturer’s instructions. Labeling and detection of apoptotic cells was completed using the Invitrogen Click-iT Plus TUNEL Assay for In Situ Apoptosis Detection 488 kit (Invitrogen, Waltham, MA, catalog #: C10617) according to manufacturer’s instructions. Nuclei were stained using DAPI (ThermoFisher Scientific, Waltham, MA, catalog #: D1306) and imaged using ZOE Fluorescent Cell Image (Bio-Rad Laboratories, Richmond, CA). Images were analyzed using FIJI. For TUNEL positive cells, threshold value was set to 14. For analyzing particles, counted those particles with size = 250-Infinity and circularity = 0.4-1.

### LNA24 transfection with leChRO-seq and RNA-seq cross comparison

HCT116 cells were plated in DMEM+10% FBS media at a density of 102,000 cells/well in a 6-well plate. Cells incubated for 24 hours in a 37 °C incubator and transfected with hsa-miR-24-3p miRCURY LNA miRNA Power Inhibitor (Qiagen, Germantown, MD, catalog #: YI04101706-DDA) or Power Negative Control A (Qiagen, Germantown, MD, catalog #: YI00199006-DDA) to a final concentration of 100 nM using Lipofectamine 3000 (ThermoFisher Scientific, Waltham, MA, catalog #: L3000-008). After 24-hours, media was replaced with complete media. After 48-hours post-transfection, cells were resuspended using 0.25% Trypsin (ThermoFisher Scientific, Waltham, MA, catalog #: 25200-114). Wells from the same treatment condition were pooled together into a single tube during each experimental replicate. 20,000 cells were isolated for total RNA isolation using the Total RNA Purification Kit (Norgen Biotek, Thorold, ON, Canada) according to manufacturer’s instructions. RNA-seq and quantitative qPCR were performed as described previously.

The remaining cells (450,000+ cells) were flash frozen using 100% EtOH and dry ice until utilized for Length Extension Chromatin Run-On Sequencing (leChRO-seq) as previously described^28,77^. <u>Chromatin Isolation:</u> Chromatin was isolated by treating cell pellet with 750µL 1X NUN buffer (20 mM HEPES, 7.5 mM MgCl2, 0.2 mM EDTA, 0.3 M NaCl, 1 M Urea, 1% NP-40, 1mM DTT, and 50 units/mL RNase Cocktail Enzyme (ThermoFisher Scientific, Waltham, MA), and 1X 50X Protease Inhibitor Cocktail (Roche, Branchburg, NJ)). Samples were vortexed vigorously for 1 minute to physically lyse the samples. An additional 750µL of 1X NUN buffer was added and samples were vortexed again for 1 minute. Cell lysates were incubated in an Eppendorf Thermomixer (Eppendorf, Hamburg, Germany) at 12°C and shaken at 2000 rpm for 30 minutes. Chromatin was pelleted by centrifuging samples at 12,500 x g for 30 minutes at 4 °C. Supernatant was removed and chromatin was washed 3 times with 1 mL 50 mM Tris-HCl (pH=7.5) containing 40 units/mL SUPERase In RNase Inhibitor. After removing supernatant from final wash, 50 µL storage buffer was added to chromatin and samples were transferred to 1.5 mL Bioruptor Microtubes with Caps for Bioruptor (Diagenode, Denville, NJ). Samples were then loaded into Pico Biorupter (Diagenode, Denville, NJ) and sonicated on high for 10 cycles (1 cycle = 30 seconds on, 30 seconds off). Sonication was repeated until chromatin was solubilized (max 3 cycles). Samples were stored at -80 °C until further processing.

#### ChRO-seq library preparation and sequencing

50 µL of 2X Biotin-11 Reaction mix (10 mM Tris-HCl pH=8.0, 5 mM MgCl2, 1 mM DTT, 300 mM KCl, 400 µM ATP, 0.8 µM CTP, 400 µM GTP, 400 µM UTP, 40 µM Biotin-11-CTP (Perkin Elmer, Waltham, MA, NEL542001EA), 100 ng yeast tRNA (VWR, Radnor, PA, 80054-306), 0.8 units/µL SUPERase In RNase Inhibitor, 1% sarkosyl) was added to 50 µL solubilized chromatin. Samples were placed in Eppendorf Thermomixer at 37°C for 5 minutes and shaken at 750 rpm. Run-on was halted by adding 300 µL Trizol LS (Life Technologies, 10296–010) and allowing the samples to incubate at room temperature for 3 minutes. RNA was purified using streptavidin beads (New England Biolabs, Ipswich, MA, S1421S) and ethanol precipitated with the co-precipitate GlycoBlue (Ambion, AM9515). Ligation of the 3’ adapter was done using the T4 RNA Ligase 1 (New England Biolabs, Ipswich, MA, M0204L). Ligation of 5’ adaptor required (i) Removal of the 5’ cap using RNA 5’ pyrophosphohydrolase (RppH, New England Biolabs, Ipswich, MA, M0356S) (ii) Phosphorylation of the 5’ end using T4 polynucleotide kinase (New England Biolabs, Ipswich, MA, M0201L) (iii) 5’ adaptor ligation using T4 RNA Ligase 1 (New England Biolabs, Ipswich, MA, M0204L). Generation of cDNA was done by using Superscript III Reverse Transcriptase (Life Technologies, 18080–044). Amplification was completed by using Q5 High-Fidelity DNA Polymerase (New England Biolabs, Ipswich, MA, M0491L). Single-end sequencing (5’ end, 75 bp) was performed at the Cornell Biotechnology Research Center using the NextSeq500 (Illumina, San Diego, CA) platform. <u>Data analysis:</u> To prepare bigwig files for further analyses, leChRO-seq libraries were aligned to the hg38 genome using the proseq2.0 pipeline (https://github.com/Danko-Lab/proseq2.0) in single-end mode (Chu et al., 2018)^28^. Annotation of leChRO-seq reads excluded reads within 500 bp downstream of the transcription start site (TSS) to account for RNA polymerase pausing at the gene promoters. Genes <1000 bp were then excluded to account for the bias resulting from short gene bodies. ChRO-seq reads were normalized and differential expression analysis was performed using DESeq2.

### Statistics

All statistical tests used are detailed in the figure legends. Either two-tailed Welch t-test (calculated using R) or two-tailed Student’s t-test (calculated using excel) was applied to datasets that were normalized (DESeq2, log2, rlog). Significance for data sets that did not statistically differ from a normal distribution (Shapiro-Wilk test p-value > 0.05) was calculated using a t-test. A two-sided Wilcoxon test was applied to non-parametric data sets unless where indicated. P-values < 0.05 are considered statistically significant. NS. = not significant, * = p<0.05, ** = p<0.01, *** = p<0.001.

## Data availability

The dataset(s) supporting the conclusions of this article are available in the Gene Expression Omnibus (GEO) repository, accession GSE188212 https://www.ncbi.nlm.nih.gov/geo/query/acc.cgi?acc=GSE188212. DESeq2 differential expression statistics are available at https://jwvillan.shinyapps.io/ME-MIRAGE/.

## Author Contributions

J.W.V, C.G.D, and P.S designed the research. L.K completed miR-24 inhibitor optimization and cellTiter-glo/cell count analyses in HCT116 following miR-24 inhibition. T.H, S.A.M, and L.E.D generated and/or expanded genetically modified mouse enteroid models. M.T.S cultured WT mouse enteroids for miR-24 inhibitor and Tgf-β experiments. M.K downloaded and generated plot of miR-24 expression across TCGA tumor types. J.W.V completed remaining wet lab experiments and computational analyses. J.W.V, C.G.D, and P.S wrote the paper. All authors reviewed and approved the paper.

## References

1. Sung, H.; Ferlay, J.; Siegel, R.L.; Laversanne, M.; Soerjomataram, I.; Jemal, A.; Bray, F. Global Cancer Statistics 2020: GLOBOCAN Estimates of Incidence and Mortality Worldwide for 36 Cancers in Countries. CA: A Cancer Journal for Clinicians 2021, 71, 209–249. doi:10.3322/caac.21660.

2. Network, T.C.G.A. Comprehensive molecular characterization of human colon and rectal cancer. Nature 2012, 487, 330 – 337. doi:10.1038/nature11252.

3. Dienstmann, R.; Wang, X.; s, A.e.l.d.R.e.; Schlicker, A.; Soneson, C.; Marisa, L.; Roepman, P.; Nyamundanda, G.; Angelino, P.; Bot, B.M.; et al. The consensus molecular subtypes of colorectal cancer. Nature Medicine 2015, pp. 1 – 13. doi:10.1038/nm.3967.

4. Molinari, C.; Marisi, G.; Passardi, A.; Matteucci, L.; Maio, G.D.; Ulivi, P. Heterogeneity in Colorectal Cancer: A Challenge for Personalized Medicine? International Journal of Molecular Sciences 2018, 19, 3733. doi:10.3390/ijms19123733.

5. Punt, C.J.A.; Koopman, M.; Vermeulen, L. From tumour heterogeneity to advances in precision treatment of colorectal cancer. Nature reviews. Clinical oncology 2017-04, 14, 235 – 246. doi:10.1038/nrclinonc.2016.171.

6. Roock, W.D.; Claes, B.; Bernasconi, D.; Schutter, J.D.; Biesmans, B.; Fountzilas, G.; Kalogeras, K.T.; Kotoula, V.; Papamichael, D.; Laurent-Puig, P.; et al. Effects of KRAS, BRAF, NRAS, and PIK3CA mutations on the efficacy of cetuximab plus chemotherapy in chemotherapy-refractory metastatic colorectal cancer: a retrospective consortium analysis. The Lancet Oncology 2010, 11, 753–762. doi:10.1016/s1470-2045(10)70130-3.

7. Sen, M.;Wang, X.; Hamdan, F.H.; Rapp, J.; Eggert, J.; Kosinsky, R.L.;Wegwitz, F.; Kutschat, A.P.; Younesi, F.S.; Gaedcke, J.; et al. ARID1A facilitates KRAS signaling-regulated enhancer activity in an AP1-dependent manner in colorectal cancer cells. Clinical Epigenetics 2019, 11, 92. doi:10.1186/s13148-019-0690-5.

8. Rahnamoun, H.; Lu, H.; Duttke, S.H.; Benner, C.; Glass, C.K.; Lauberth, S.M. Mutant p53 shapes the enhancer landscape of cancer cells in response to chronic immune signaling. Nature Communications 2017, pp. 1 – 14. doi:10.1038/s41467-017-01117-y.

9. Necela, B.M.; Carr, J.M.; Asmann, Y.W.; Thompson, E.A. Differential Expression of MicroRNAs in Tumors from Chronically Inflamed or Genetic (APCMin/+) Models of Colon Cancer. PLoS ONE 2011, 6, e18501. doi:10.1371/journal.pone.0018501.

10. Bailey, J.M.; Hendley, A.M.; Lafaro, K.J.; Pruski, M.A.; Jones, N.C.; Alsina, J.; Younes, M.; Maitra, A.; McAllister, F.; Iacobuzio-Donahue, C.A.; et al. p53 mutations cooperate with oncogenic Kras to promote adenocarcinoma from pancreatic ductal cells. Oncogene 2016, 35, 4282 – 4288. doi:10.1038/onc.2015.441.

11. Trobridge, P.; Knoblaugh, S.; Washington, M.K.; Munoz, N.M.; Tsuchiya, K.D.; Rojas, A.; Song, X.; Ulrich, C.M.; Sasazuki, T.; Shirasawa, S.; et al. TGF-beta receptor inactivation and mutant Kras induce intestinal neoplasms in mice via a beta-catenin-independent pathway. Gastroenterology 2009-05, 136, 1680 8.e7. doi:10.1053/j.gastro.2009.01.066.

12. Han, T.; Goswami, S.; Hu, Y.; Tang, F.; Zafra, M.P.; Murphy, C.; Cao, Z.; Poirier, J.T.; Khurana, E.; Elemento, O.; et al. Lineage reversion drives WNT independence in intestinal cancer. Cancer discovery 2020, pp. CD–19–1536. doi:10.1158/2159-8290.cd-19-1536.

13. Liu, F.; Hon, G.C.; Villa, G.R.; Turner, K.M.; Ikegami, S.; Yang, H.; Ye, Z.; Li, B.; Kuan, S.; Lee, A.Y.; et al. EGFR Mutation Promotes Glioblastoma through Epigenome and Transcription Factor Network Remodeling. Molecular Cell 2015, 60, 307 – 318. doi:10.1016/j.molcel.2015.09.002.

14. Ibrahim, H.; Lim, Y.C. KRAS-associated microRNAs in colorectal cancer. Oncology Reviews 2020, 14, 454. doi:10.4081/oncol.2020.454.

15. Brown, D.; Rahman, M.; Nana-Sinkam, S.P. MicroRNAs in Respiratory Disease. A Clinician’s Overview. Annals of the American Thoracic Society 2014, 11, 1277–1285. doi:10.1513/annalsats.201404-179fr.

16. Fridrichova, I.; Zmetakova, I. MicroRNAs Contribute to Breast Cancer Invasiveness. Cells 2019, 8, 1361. doi:10.3390/cells8111361.

17. Hao, N.B.; He, Y.F.; Li, X.Q.; Wang, K.; Wang, R.L. The role of miRNA and lncRNA in gastric cancer. Oncotarget 2015, 5, 81572–81582. doi:10.18632/oncotarget.19197.

18. Ma, L.; Reinhardt, F.; Pan, E.; Soutschek, J.; Bhat, B.; Marcusson, E.; Teruya-Feldstein, J.; Bell, G.W.; Weinberg, R.A. Therapeutic silencing of miR-10b inhibits metastasis in a mouse mammary tumor model. Nature biotechnology 2010, 28, 341–347. doi:10.1038/nbt.1618.

19. Teplyuk, N.M.; Uhlmann, E.J.; Gabriely, G.; Volfovsky, N.;Wang, Y.; Teng, J.; Karmali, P.; Marcusson, E.; Peter, M.; Mohan, A.; et al. Therapeutic potential of targeting microRNA-10b in established intracranial glioblastoma: first steps toward the clinic. EMBO molecular medicine 2016, 8, 268–87. doi:10.15252/emmm.201505495.

20. Hanna, J.; Hossain, G.S.; Kocerha, J. The Potential for microRNA Therapeutics and Clinical Research. Frontiers in Genetics 2019, 10, 478. doi:10.3389/fgene.2019.00478.

21. Seto, A.G.; Beatty, X.; Lynch, J.M.; Hermreck, M.; Tetzlaff, M.; Duvic, M.; Jackson, A.L. Cobomarsen, an oligonucleotide inhibitor of miR-155, co-ordinately regulates multiple survival pathways to reduce cellular proliferation and survival in cutaneous T-cell lymphoma. British Journal of Haematology 2018, 183, 428– 444. doi:10.1111/bjh.15547.

22. Cojocneanu, R.; Braicu, C.; Raduly, L.; Jurj, A.; Zanoaga, O.; Magdo, L.; Irimie, A.; Muresan, M.S.; Ionescu, C.; Grigorescu, M.; et al. Plasma and Tissue Specific miRNA Expression Pattern and Functional Analysis Associated to Colorectal Cancer Patients. Cancers 2020, 12, 843. doi:10.3390/cancers12040843.

23. Falzone, L.; Scola, L.; Zanghì, A.; Biondi, A.; Cataldo, A.D.; Libra, M.; Candido, S. Integrated analysis of colorectal cancer microRNA datasets: identification of microRNAs associated with tumor development. Aging (Albany NY) 2018, 10, 1000–1014. doi:10.18632/aging.101444.

24. Bandrés, E.; Cubedo, E.; Agirre, X.; Malumbres, R.; Zárate, R.; Ramirez, N.; Abajo, A.; Navarro, A.; Moreno, I.; Monzó, M.; et al. Identification by Real-time PCR of 13 mature microRNAs differentially expressed in colorectal cancer and non-tumoral tissues. Molecular Cancer 2006, 5, 29–29. doi:10.1186/1476-4598-5-29.

25. To, K.K.; Tong, C.W.;Wu, M.; Cho,W.C. MicroRNAs in the prognosis and therapy of colorectal cancer: From bench to bedside. World Journal of Gastroenterology 2018, 24, 2949–2973. doi:10.3748/wjg.v24.i27.2949.

26. Berg, K.C.G.; Eide, P.W.; Eilertsen, I.A.; Johannessen, B.; Bruun, J.; Danielsen, S.A.; Bjørnslett, M.; Meza-Zepeda, L.A.; Eknæs, M.; Lind, G.E.; et al. Multi-omics of 34 colorectal cancer cell lines - a resource for biomedical studies Molecular Cancer 2017. pp. 1 – 16. doi:10.1186/s12943-017-0691-y.

27. Crespo, M.; Vilar, E.; Tsai, S.Y.; Chang, K.; Amin, S.; Srinivasan, T.; Zhang, T.; Pipalia, N.H.; Chen, H.J.;Witherspoon, M.; et al. Colonic organoids derived from human induced pluripotent stem cells for modeling colorectal cancer and drug testing. Nature Medicine 2017, 23, 878 – 884. doi:10.1038/nm.4355.

28. Chu, T.; Rice, E.J.; Booth, G.T.; Salamanca, H.H.;Wang, Z.; Core, L.J.; Longo, S.L.; Corona, R.J.; Chin, L.S.; Lis, J.T.; et al. Chromatin run-on and sequencing maps the transcriptional regulatory landscape of glioblastoma multiforme. Nature Genetics 2018-11, 50, 1553 – 1564. doi:10.1038/s41588-018-0244-3.

29. Schatoff, E.M.; Goswami, S.; Zafra, M.P.; Foronda, M.; Shusterman, M.; Leach, B.I.; Katti, A.; Diaz, B.J.; Dow, L.E. Distinct CRC-associated APC mutations dictate response to Tankyrase inhibition. Cancer discovery 2019, pp. CD–19–0289. doi:10.1158/2159-8290.cd-19-0289.

30. Dow, L.E.; O’Rourke, K.P.; Simon, J.; Tschaharganeh, D.F.; Es, J.H.v.; Clevers, H.; Lowe, S.W. Apc Restoration Promotes Cellular Differentiation and Reestablishes Crypt Homeostasis in Colorectal Cancer. Cell 2015, 161, 1539 – 1552. doi:10.1016/j.cell.2015.05.033.

31. Han, T.; Schatoff, E.M.; Murphy, C.; Zafra, M.P.; Wilkinson, J.E.; Elemento, O.; Dow, L.E. R-Spondin chromosome rearrangements drive Wnt-dependent tumour initiation and maintenance in the intestine. Nature Communications 2017, 8, 1 – 12. doi:10.1038/ncomms15945.

32. Kanke, M.; Baran-Gale, J.; Villanueva, J.; Sethupathy, P. miRquant 2.0: an Expanded Tool for Accurate Annotation and Quantification of MicroRNAs and their isomiRs from Small RNA-Sequencing Data. Journal of integrative bioinformatics 2016, 13, 307. doi:10.2390/biecoll-jib-2016-307.

33. Pantano, L. DEG Report: Report of DEG analysis. doi:10.18129/B9.bioc.DEGreport.

34. Lu, D.; Yao, Q.; Zhan, C.; Le-Meng, Z.; Liu, H.; Cai, Y.; Tu, C.; Li, X.; Zou, Y.; Zhang, S. MicroRNA-146a promote cell migration and invasion in human colorectal cancer via carboxypeptidase M/src-FAK pathway. Oncotarget 2017, 8, 22674–22684. doi:10.18632/oncotarget.15158.

35. Khorrami, S.; Hosseini, A.Z.; Mowla, S.J.; Soleimani, M.; Rakhshani, N.; Malekzadeh, R. MicroRNA-146a induces immune suppression and drug-resistant colorectal cancer cells. Tumor Biology 2017, 39, 1010428317698365. doi:10.1177/1010428317698365.

36. Kawaguchi, Y.; Kopetz, S.; Newhook, T.E.; Bellis, M.D.; Chun, Y.S.; Tzeng, C.W.D.; Aloia, T.A.; Vauthey, J.N. Mutation Status of RAS, TP53, and SMAD4 is Superior to Mutation Status of RAS Alone for Predicting Prognosis after Resection of Colorectal Liver Metastases. Clinical Cancer Research 2019, 25, 5843–5851. doi:10.1158/1078-0432.ccr-19-0863.

37. Matano, M.; Date, S.; Shimokawa, M.; Takano, A.; Fujii, M.; Ohta, Y.;Watanabe, T.; Kanai, T.; Sato, T. Modeling colorectal cancer using CRISPR-Cas9–mediated engineering of human intestinal organoids. Nature Medicine 2015, 21, 256–262. doi:10.1038/nm.3802.

38. Xie, M.; Qin, H.; Luo, Q.; Huang, Q.; He, X.; Yang, Z.; Lan, P.; Lian, L. MicroRNA-30a regulates cell proliferation and tumor growth of colorectal cancer by targeting CD73. BMC Cancer 2017, 17, 305. doi:10.1186/s12885-017-3291-8.

39. Baraniskin, A.; Birkenkamp-Demtroder, K.; Maghnouj, A.; Zöllner, H.; Munding, J.; Klein-Scory, S.; Reinacher-Schick, A.; Schwarte-Waldhoff, I.; Schmiegel,W.; Hahn, S.A. MiR-30a-5p suppresses tumor growth in colon carcinoma by targeting DTL. Carcinogenesis 2012, 33, 732–739. doi:10.1093/carcin/bgs020.

40. Liang, Z.; Li, X.; Liu, S.; Li, C.; Wang, X.; Xing, J. MiR-141–3p inhibits cell proliferation, migration and invasion by targeting TRAF5 in colorectal cancer. Biochemical and Biophysical Research Communications 2019, 514, 699–705. doi:10.1016/j.bbrc.2019.05.002.

41. Long, Z.H.; Bai, Z.G.; Song, J.N.; Zheng, Z.; Li, J.; Zhang, J.; Cai, J.; Yao, H.W.; Wang, J.; Yang, Y.C.; et al. miR-141 Inhibits Proliferation and Migration of Colorectal Cancer SW480 Cells. Anticancer Research 2017, 37, 4345–4352. doi:10.21873/anticanres.11828.

42. Li, Y.; Lauriola, M.; Kim, D.; Francesconi, M.; D’Uva, G.; Shibata, D.; Malafa, M.P.; Yeatman, T.J.; Coppola, D.; Solmi, R.; et al. Adenomatous polyposis coli (APC) regulates miR17-92 cluster through - catenin pathway in colorectal cancer. Oncogene 2016, 35, 4558 – 4568. doi:10.1038/onc.2015.522.

43. Guo, X.; Zhu, Y.; Hong, X.; Zhang, M.; Qiu, X.; Wang, Z.; Qi, Z.; Hong, X. miR-181d and c-myc-mediated inhibition of CRY2 and FBXL3 reprograms metabolism in colorectal cancer. Cell death & disease 2016, 8, e2958. doi:10.1038/cddis.2017.300.

44. Yamazaki, N.; Koga, Y.; Taniguchi, H.; Kojima, M.; Kanemitsu, Y.; Saito, N.; Matsumura, Y. High expression of miR-181c as a predictive marker of recurrence in stage II colorectal cancer. Oncotarget 2016, 8, 6970–6983. doi:10.18632/oncotarget.14344.

45. Pan, S.; Deng, Y.; Fu, J.; Zhang, Y.; Zhang, Z.; Qin, X. N6-methyladenosine upregulates miR-181d-5p in exosomes derived from cancer-associated fibroblasts to inhibit 5-FU sensitivity by targeting NCALD in colorectal cancer. International Journal of Oncology 2022, 60, 14. doi:10.3892/ijo.2022.5304.

46. Dinh, T.A.; Jewell, M.L.; Kanke, M.; Francisco, A.; Sritharan, R.; Turnham, R.E.; Lee, S.; Kastenhuber, E.R.; Wauthier, E.; Guy, C.D.; et al. MicroRNA-375 Suppresses the Growth and Invasion of Fibrolamellar Carcinoma. Cellular and molecular gastroenterology and hepatology 2019, 7, 803 – 817. doi:10.1016/j.jcmgh.2019.01.008.

47. Kang, W.; Huang, T.; Zhou, Y.; Zhang, J.; Lung, R.W.M.; Tong, J.H.M.; Chan, A.W.H.; Zhang, B.; Wong, C.C.; Wu, F.; et al. miR-375 is involved in Hippo pathway by targeting YAP1/TEAD4-CTGF axis in gastric carcinogenesis. Cell Death & Disease 2018, 9, 92. doi:10.1038/s41419-017-0134-0.

48. Alam, K.J.; Mo, J.S.; Han, S.H.; Park, W.C.; Kim, H.S.; Yun, K.J.; Chae, S.C. MicroRNA 375 regulates proliferation and migration of colon cancer cells by suppressing the CTGF-EGFR signaling pathway. International Journal of Cancer 2017, 141, 1614 – 1629. doi:10.1002/ijc.30861.

49. Yun, S.M.; Kim, S.H.; Kim, E.H. The Molecular Mechanism of Transforming Growth Factor-Signaling for Intestinal Fibrosis: A Mini-Review. Frontiers in pharmacology 2019, 10, 162. doi:10.3389/fphar.2019.00162.

50. Love, M.I.; Huber, W.; Anders, S. Moderated estimation of fold change and dispersion for RNA-seq data with DESeq2. Genome biology 2014, 15, 550. doi:10.1186/s13059-014-0550-8.

51. Baran-Gale, J.; Fannin, E.E.; Kurtz, C.L.; Sethupathy, P. Beta cell 5’-shifted isomiRs are candidate regulatory hubs in type 2 diabetes. PLOS ONE 2013, 8, e73240. doi:10.1371/journal.pone.0073240.

52. Chen, E.Y.; Tan, C.M.; Kou, Y.; Duan, Q.; Wang, Z.; Meirelles, G.V.; Clark, N.R.; Ma’ayan, A. Enrichr: interactive and collaborative HTML5 gene list enrichment analysis tool. BMC Bioinformatics 2013, 14, 128–128. doi:10.1186/1471-2105-14-128.

53. Kuleshov, M.V.; Jones, M.R.; Rouillard, A.D.; Fernandez, N.F.; Duan, Q.; Wang, Z.; Koplev, S.; Jenkins, S.L.; Jagodnik, K.M.; Lachmann, A.; et al. Enrichr: a comprehensive gene set enrichment analysis web server 2016 update. Nucleic Acids Research 2016, 44, W90–W97. doi:10.1093/nar/gkw377.

54. Xie, Z.; Bailey, A.; Kuleshov, M.V.; Clarke, D.J.B.; Evangelista, J.E.; Jenkins, S.L.; Lachmann, A.; Wojciechowicz, M.L.; Kropiwnicki, E.; Jagodnik, K.M.; et al. Gene Set Knowledge Discovery with Enrichr. Current Protocols 2021, 1, e90. doi:10.1002/cpz1.90.

55. Sarma, S.N.; Kim, Y.J.; Song, M.; Ryu, J.C. Induction of apoptosis in human leukemia cells through the production of reactive oxygen species and activation of HMOX1 and Noxa by benzene, toluene, and oxylene. Toxicology 2011, 280, 109–117. doi:10.1016/j.tox.2010.11.017.

56. Kwon, M.Y.; Park, E.; Lee, S.J.; Chung, S.W. Heme oxygenase-1 accelerates erastin-induced ferroptotic cell death. Oncotarget 2015, 6, 24393 – 24403. doi:10.18632/oncotarget.5162.

57. Zhang, L.; Jia, G.; Shi, B.; Ge, G.; Duan, H.; Yang, Y. PRSS8 is Downregulated and Suppresses Tumour Growth and Metastases in Hepatocellular Carcinoma. Cellular Physiology and Biochemistry 2016, 40, 757– 769. doi:10.1159/000453136.

58. Wu, L.; Gong, Y.; Yan, T.; Zhang, H. LINP1 promotes the progression of cervical cancer by scaffolding EZH2, LSD1, and DNMT1 to inhibit the expression of KLF2 and PRSS8. Biochemistry and Cell Biology 2020, 98, 591–599. doi:10.1139/bcb-2019-0446.

59. Yang, Y.z.; Zhao, X.j.; Xu, H.j.;Wang, S.c.; Pan, Y.;Wang, S.j.; Xu, Q.; Jiao, R.q.; Gu, H.m.; Kong, L.d. Magnesium isoglycyrrhizinate ameliorates high fructose-induced liver fibrosis in rat by increasing miR-375-3p to suppress JAK2/STAT3 pathway and TGF-1/Smad signaling. Acta Pharmacologica Sinica 2019, 40, 879–894. doi:10.1038/s41401-018-0194-4.

60. Zhang, X.; Chen, Q.; Song, H.; Jiang,W.; Xie, S.; Huang, J.; Kang, G. MicroRNA-375 prevents TGF--dependent transdifferentiation of lung fibroblasts via the MAP2K6/P38 pathway. Molecular Medicine Reports 2020, 22, 1803–1810. doi:10.3892/mmr.2020.11261.

61. Wei, R.; Yang, Q.; Han, B.; Li, Y.; Yao, K.; Yang, X.; Chen, Z.; Yang, S.; Zhou, J.; Li, M.; et al. microRNA-375 inhibits colorectal cancer cells proliferation by downregulating JAK2/STAT3 and MAP3K8/ERK signaling pathways. Oncotarget 2017, 8, 16633–16641. doi:10.18632/oncotarget.15114.

62. He, X.X.; Chang, Y.; Meng, F.Y.; Wang, M.Y.; Xie, Q.H.; Tang, F.; Li, P.Y.; Song, Y.H.; Lin, J.S. MicroRNA-375 targets AEG-1 in hepatocellular carcinoma and suppresses liver cancer cell growth in vitro and in vivo. Oncogene 2012, 31, 3357 – 3369. doi:10.1038/onc.2011.500.

63. Peck, B.C.E.; Mah, A.T.; Pitman, W.A.; Ding, S.; Lund, P.K.; Sethupathy, P. Functional Transcriptomics in Diverse Intestinal Epithelial Cell Types Reveals Robust MicroRNA Sensitivity in Intestinal Stem Cells to Microbial Status. Journal of Biological Chemistry 2017, 292, 2586 – 2600. doi:10.1074/jbc.m116.770099.

64. Gao, Z.; Zhou, L.; Hua, S.; Wu, H.; Luo, L.; Li, L.; Wang, S.; Liu, Y.; Zhou, Z.; Chen, X. miR-24-3p promotes colon cancer progression by targeting ING1. Signal transduction and targeted therapy 2020, 5, 171 – 3. doi:10.1038/s41392-020-0206-y.

65. Zhang, H.W.; Shi, Y.; Liu, J.B.;Wang, H.M.;Wang, P.Y.;Wu, Z.J.; Li, L.; Gu, L.P.; Cao, P.S.;Wang, G.R.; et al. Cancer-associated fibroblast-derived exosomal microRNA-24-3p enhances colon cancer cell resistance to MTX by down-regulating CDX2/HEPH axis. Journal of cellular and molecular medicine 2021. doi:10.1111/jcmm.15765.

66. Gao, Y.; Liu, Y.; Du, L.; Li, J.; Qu, A.; Zhang, X.;Wang, L.;Wang, C. Down-regulation of miR-24-3p in colorectal cancer is associated with malignant behavior. Medical oncology (Northwood, London, England) 2015-01, 32, 362 – 8. doi:10.1007/s12032-014-0362-4.

67. Zhang, Q.; Li, W.; Liu, G.; Tang, W. MicroRNA-24 regulates the growth and chemosensitivity of the human colorectal cancer cells by targeting RNA-binding protein DND1. Journal of B.U.ON. : official journal of the Balkan Union of Oncology 2019-07, 24, 1476 – 1481.

68. Chhabra, R.; Dubey, R.; Saini, N. Cooperative and individualistic functions of the microRNAs in the miR-23a27a24-2 cluster and its implication in human diseases. Molecular Cancer 2010, 9, 232–232. doi:10.1186/1476-4598-9-232.

69. Zhu, X.F.; Li,W.; Ma, J.Y.; Shao, N.; Zhang, Y.J.; Liu, R.M.; Wu,W.B.; Lin, Y.;Wang, S.M. Knockdown of heme oxygenase-1 promotes apoptosis and autophagy and enhances the cytotoxicity of doxorubicin in breast cancer cells. Oncology Letters 2015, 10, 2974–2980. doi:10.3892/ol.2015.3735.

70. Petrache, I.; Otterbein, L.E.; Alam, J.; Wiegand, G.W.; Choi, A.M.K. Heme oxygenase-1 inhibits TNF--induced apoptosis in cultured fibroblasts. American Journal of Physiology-Lung Cellular and Molecular Physiology 2000, 278, L312–L319. doi:10.1152/ajplung.2000.278.2.l312.

71. Becker, J.C.; Fukui, H.; Imai, Y.; Sekikawa, A.; Kimura, T.; Yamagishi, H.; Yoshitake, N.; Pohle, T.; Domschke, W.; Fujimori, T. Colonic expression of heme oxygenase-1 is associated with a better long-term survival in patients with colorectal cancer. Scandinavian Journal of Gastroenterology 2009, 42, 852–858. doi:10.1080/00365520701192383.

72. Ishikawa, T.; Yoshida, N.; Higashihara, H.; Inoue, M.; Uchiyama, K.; Takagi, T.; Handa, O.; Kokura, S.; Naito, Y.; Okanoue, T.; et al. Different effects of constitutive nitric oxide synthase and heme oxygenase on pulmonary or liver metastasis of colon cancer in mice. Clinical & Experimental Metastasis 2003, 20, 445– 450. doi:10.1023/a:1025448403124.

73. Hofmans, S.; Berghe, T.V.; Devisscher, L.; Hassannia, B.; Lyssens, S.; Joossens, J.; Veken, P.V.D.; Vandenabeele, P.; Augustyns, K. Novel Ferroptosis Inhibitors with Improved Potency and ADME Properties. Journal of Medicinal Chemistry 2016, 59, 2041–2053. doi:10.1021/acs.jmedchem.5b01641.

74. Guo, J.; Xu, B.; Han, Q.; Zhou, H.; Xia, Y.; Gong, C.; Dai, X.; Li, Z.;Wu, G. Ferroptosis: A Novel Antitumor Action for Cisplatin. Cancer research and treatment : official journal of Korean Cancer Association 2018-04, 50, 445 – 460. doi:10.4143/crt.2016.572.

75. Jackson, E.L.; Willis, N.; Mercer, K.; Bronson, R.T.; Crowley, D.; Montoya, R.; Jacks, T.; Tuveson, D.A. Analysis of lung tumor initiation and progression using conditional expression of oncogenic K-ras. Genes & Development 2001, 15, 3243–3248. doi:10.1101/gad.943001.

76. Grimson, A.; Farh, K.K.H.; Johnston, W.K.; Garrett-Engele, P.; Lim, L.P.; Bartel, D.P. MicroRNA Targeting Specificity in Mammals: Determinants beyond Seed Pairing. Molecular Cell 2007, 27, 91–105. doi:10.1016/j.molcel.2007.06.017.

77. Hung, Y.H.; Huang, S.; Dame, M.K.; Yu, Q.; Yu, Q.C.; Zeng, Y.A.; Camp, J.G.; Spence, J.R.; Sethupathy, P. Chromatin regulatory dynamics of early human small intestinal development using a directed differentiation model. Nucleic Acids Research 2021, 49, gkaa1204.#x2013;. doi:10.1093/nar/gkaa1204.

