## Supplemental for "Genome edited colorectal cancer organoid models reveal distinct microRNA activity patterns across different mutation profiles"

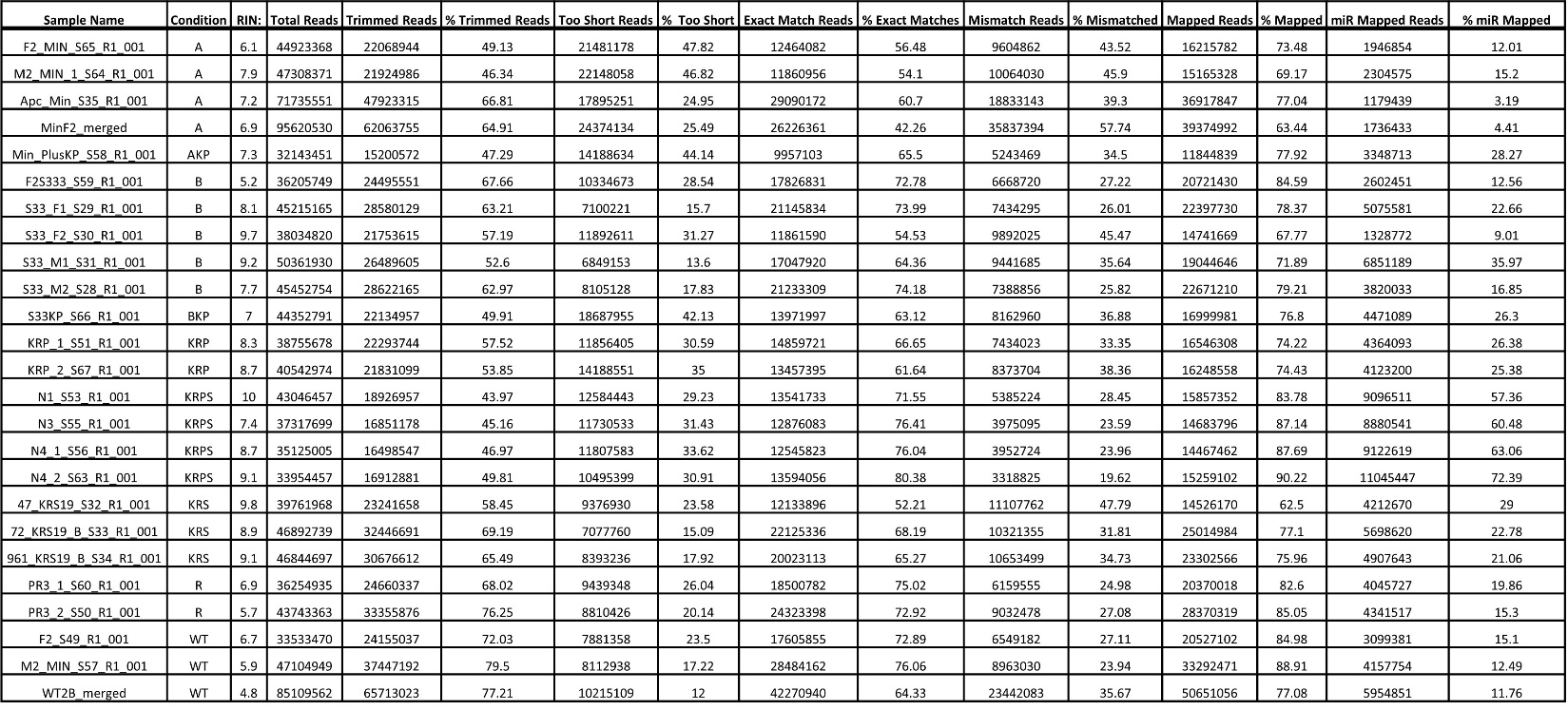
**Table S1:** Mapping statistics of small RNA-seq data produced from genetically modified mouse enteroids and WT controls.

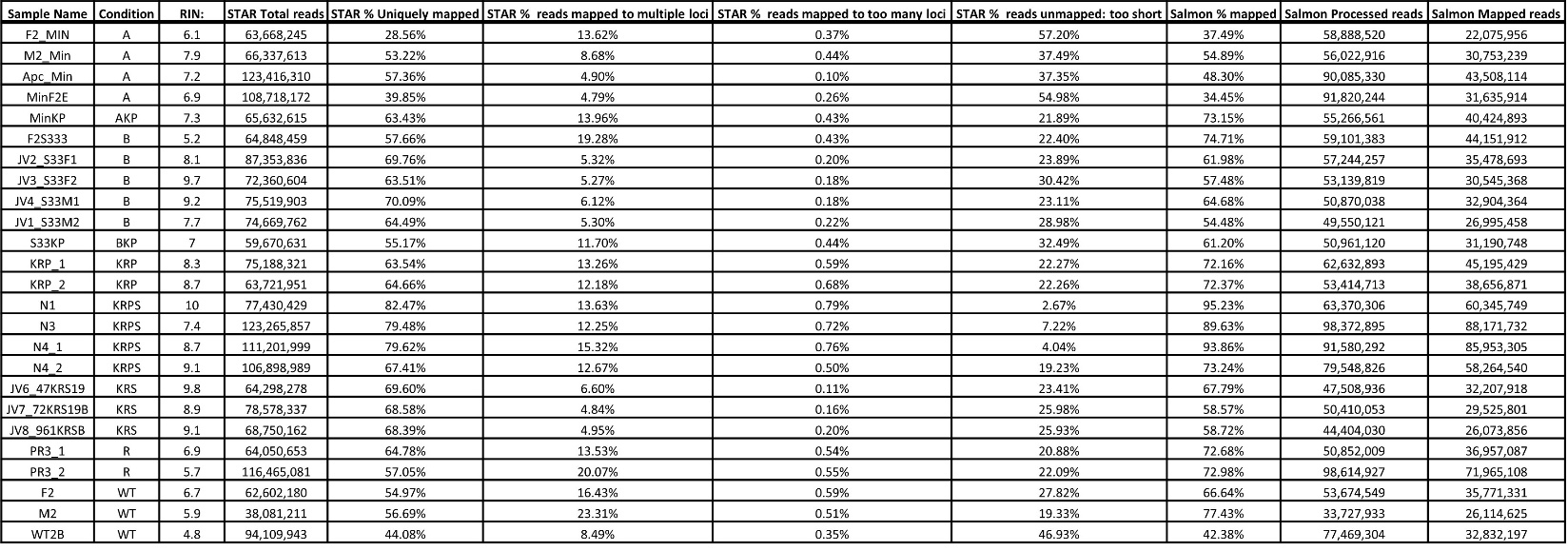

**Table S2:** Mapping statistics of RNA-seq data produced from genetically modified mouse enteroids and WT controls.

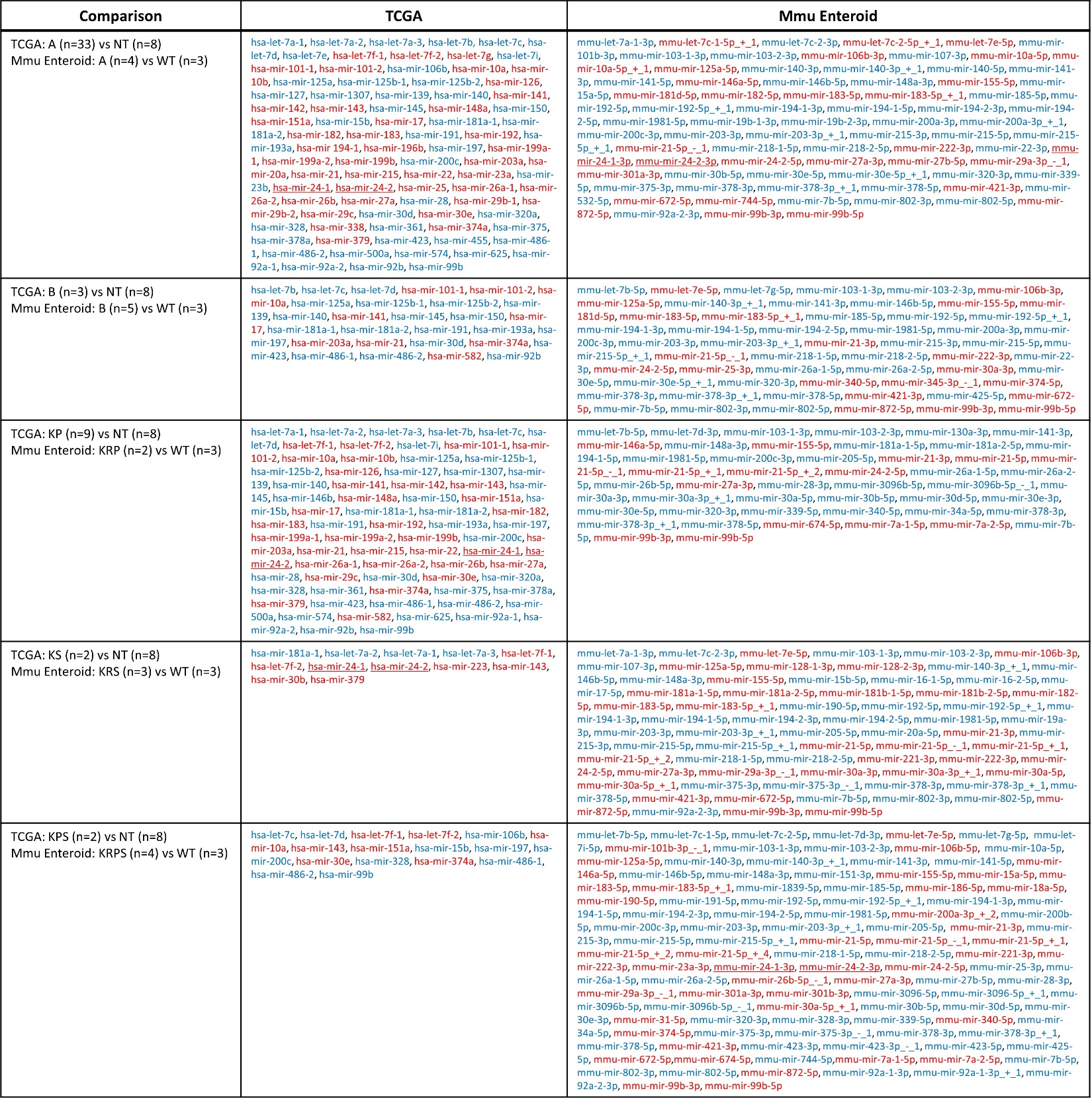

**Table S3:** Differentially expressed miRNAs when comparing TCGA tumors with specific genotypes to non-tumor (NT) controls (>500 RPMMM in either condition, p-value<0.05, hochberg p-adj<0.2, and fold change >1.5x) and mutant mouse enteroids with WT control (baseMean>500 DESeq normalized counts, p-value<0.05, DESeq2 p-adj<0.2, and fold change >1.5x). MiRNAs in red are upregulated. MiRNAs in blue are downregulated

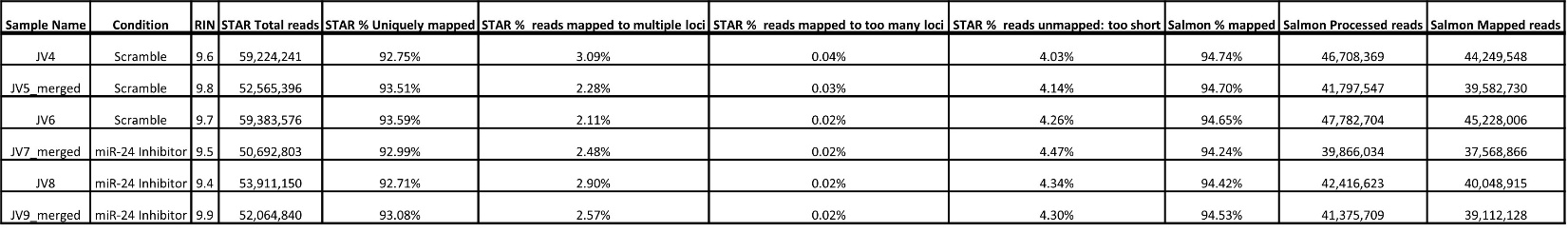

**Table S4:** Mapping statistics of RNA-seq data produced from HCT116 cells transfected with scramble control or miR-24 inhibitor.

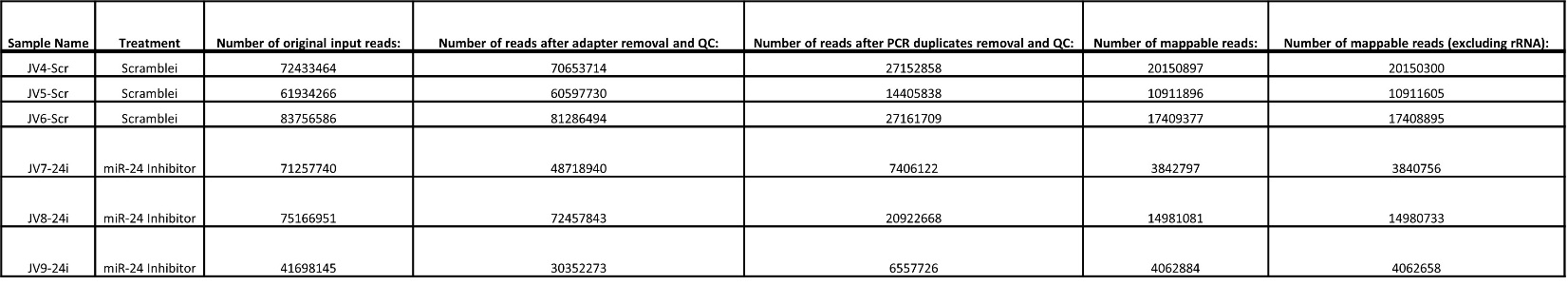

**Table S5:** Mapping statistics of ChRO-seq data produced from HCT116 cells transfected with scramble control or miR-24 inhibitor.

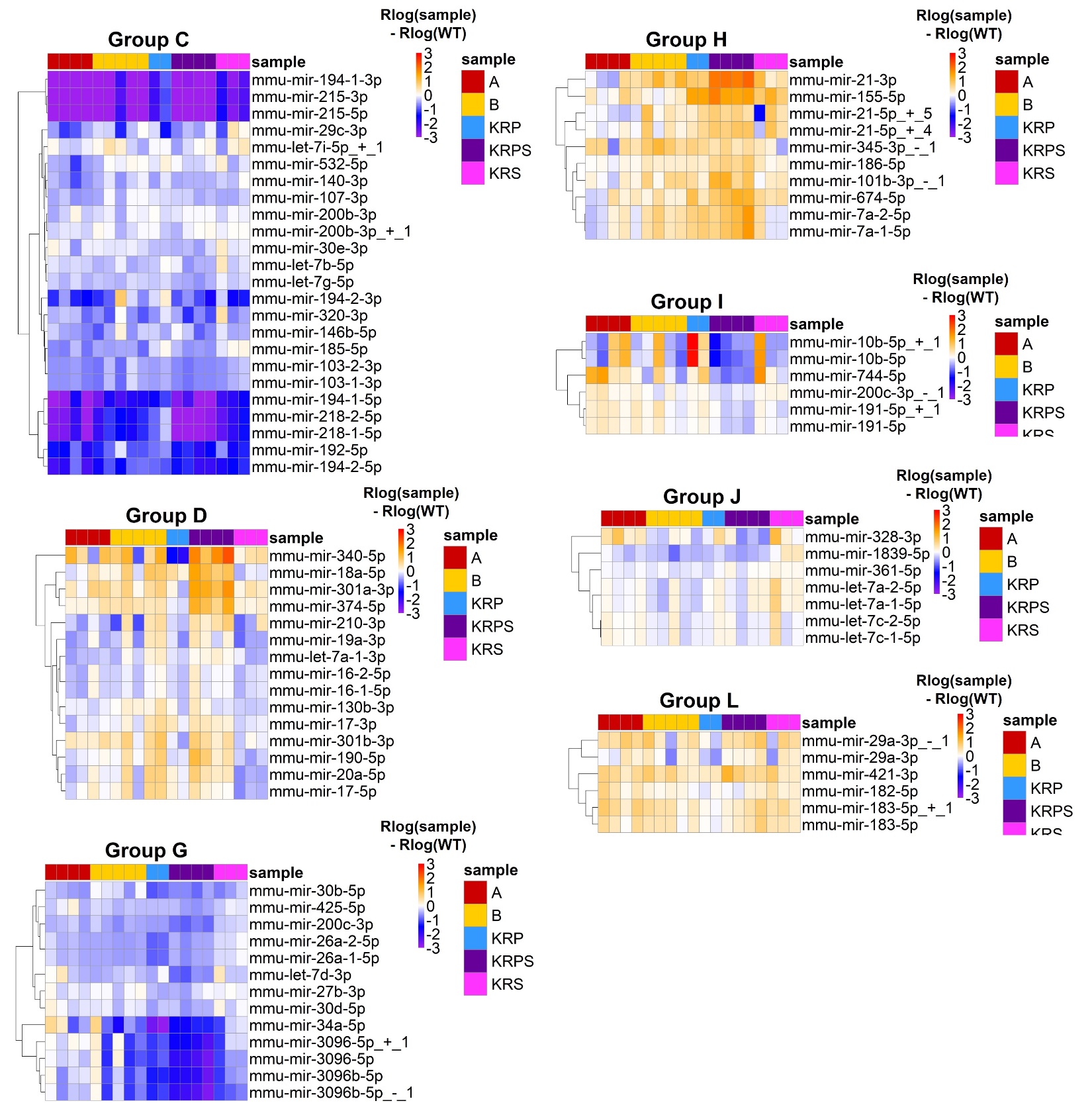

**Supplemental Figure 1:** Heatmaps show the magnitude of change in miRNA expression relative to WT by subtracting rlog normalized miRNA expression for each enteroid sample by the rlog average WT expression. Groups not shown in the main text shown here. Color intensity rlog normalized miRNA expression in each genetically modified enteroid sample subtracted from average WT. Color scale minimum saturates at -3 and maximum saturates at 3.

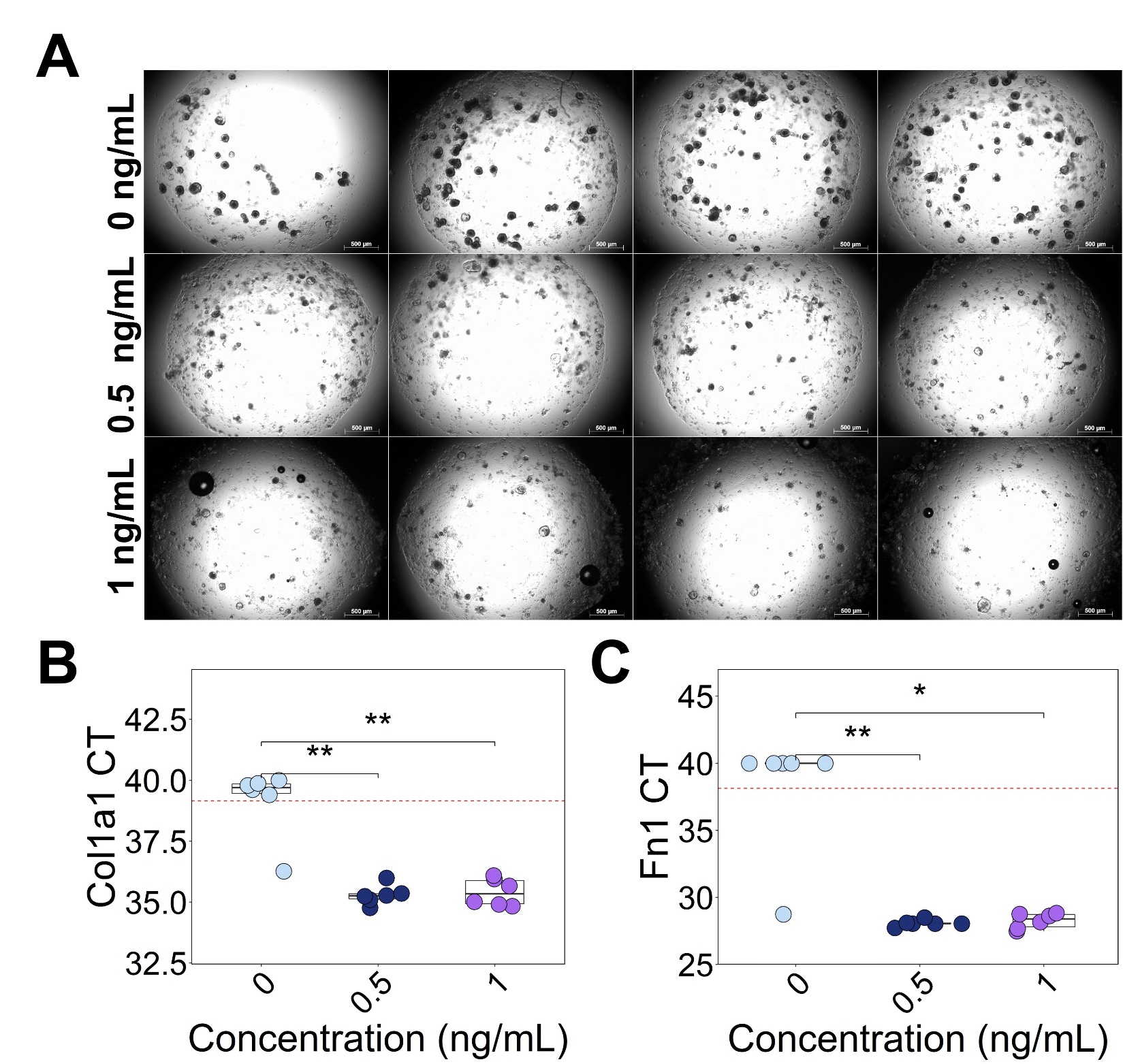

**Supplemental Figure 2:** (**A**) Brightfield images of mouse enteroids treated with 0, 0.5, or 1 ng/mL recombinant human TGF-B1. (**B**) *Col1a1* and (**C**) *Fn1* CTs from RT-qPCR. In cases for which gene expression was not detected at 40 cycles, CT was set to 40 for analysis. Significance determined by two-sided Wilcoxon test. *p<0.05, **p<0.01, ***p<0.001.

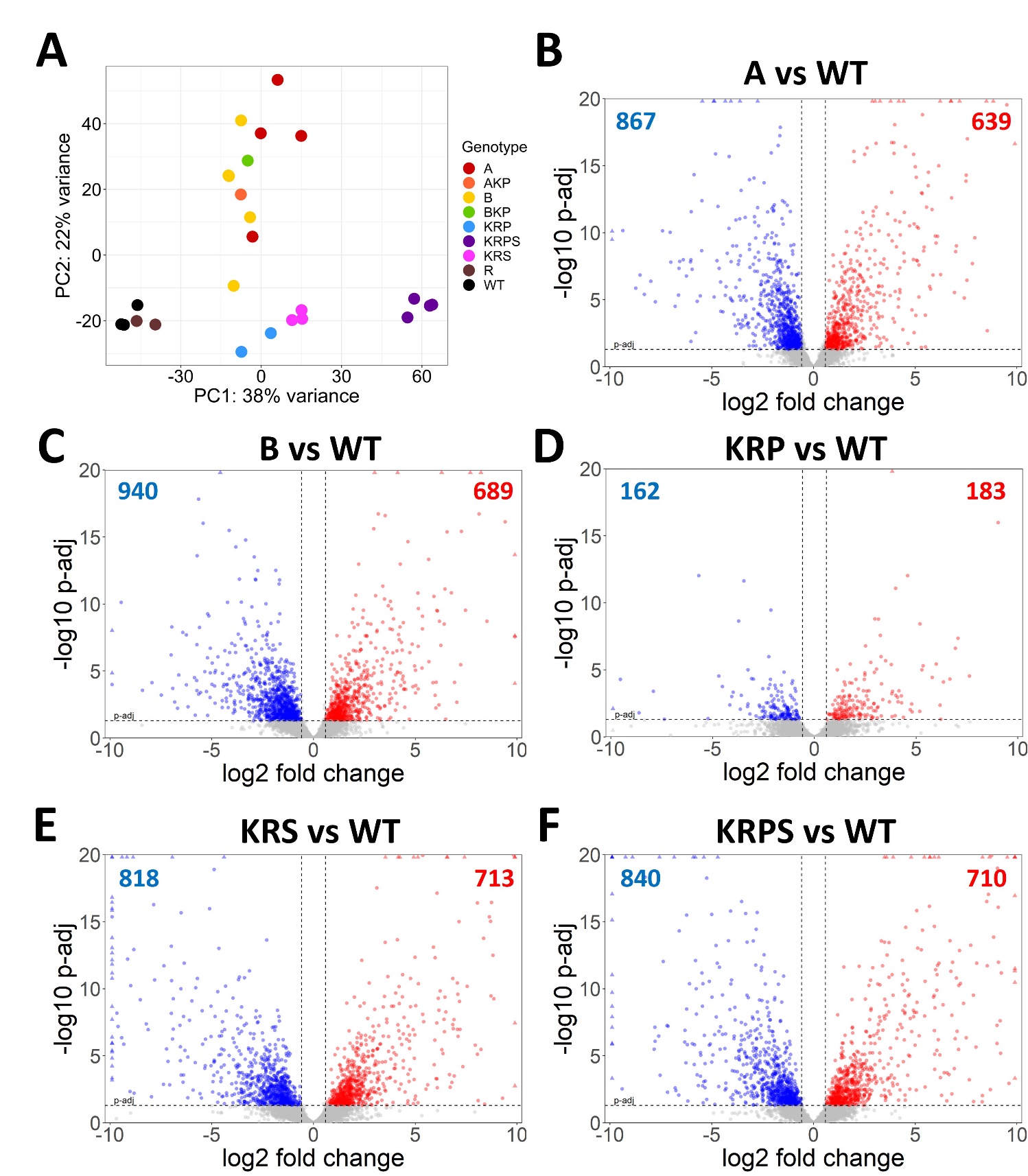
**Supplemental Figure 3:** (**A**) PCA plot generated using gene expression profiles generated from mutant mouse enteroids with wild-type controls. (**B**-**F**) Volcano plots highlighting differentially expressed genes in mutant mouse enteroids relative to WT control. Genes filtered for expression above 500 normalized counts in either condition. Horizontal dashed line represents p-adj cutoff of 0.05 (DESeq2). Vertical dashed lines represent 1.5x fold change.

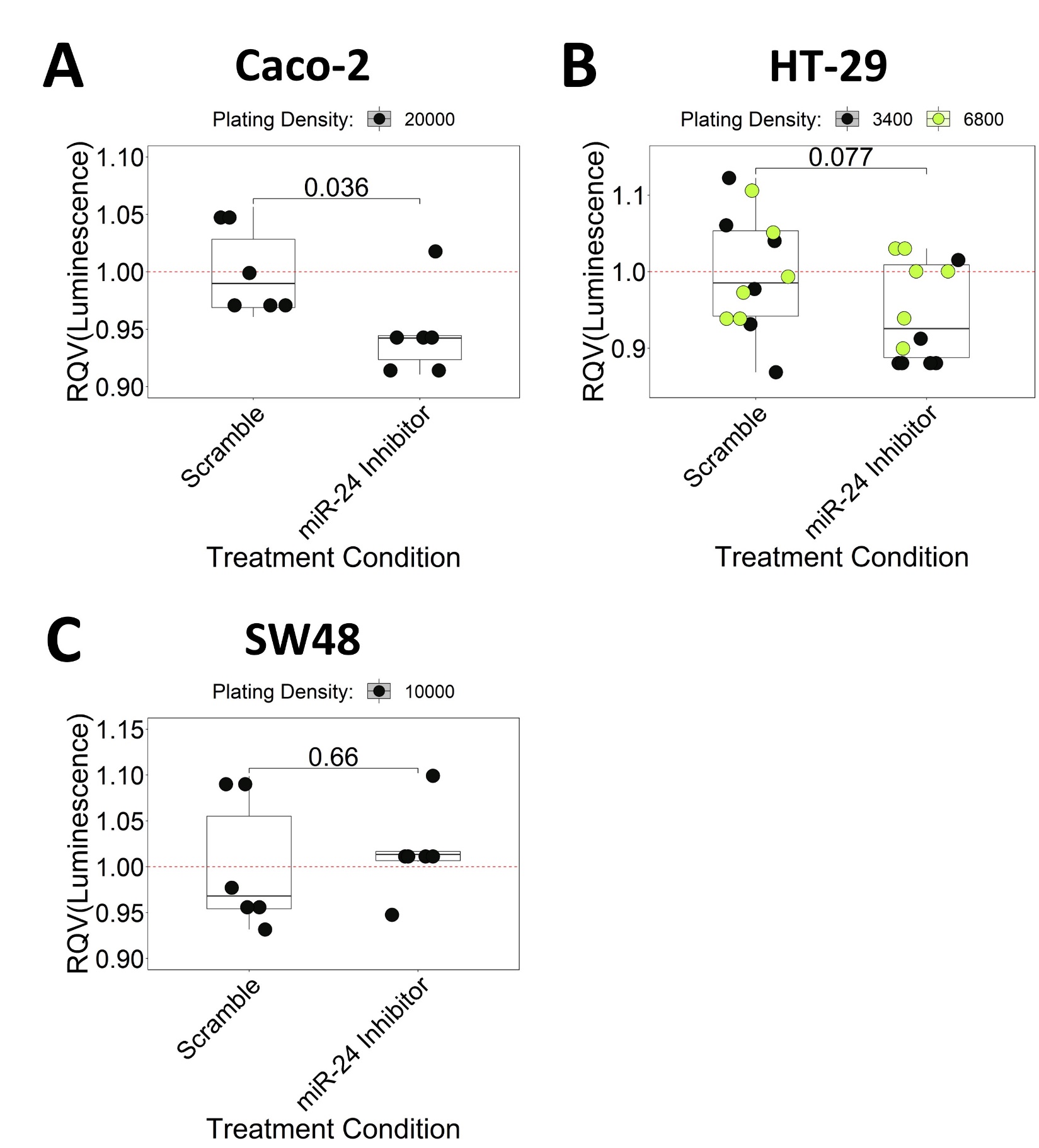

**Supplemental Figure 4:** Relative luminescent signal performed by the CellTiter-Glo assay after miR-24-3p inhibition in (**A**) Caco-2, (**B**) HT-29, and (**C**) SW48 colorectal cancer cell lines. Coloration represents the cell plating density in 96-well plates. Significance determined by two-tailed Welch t-test.

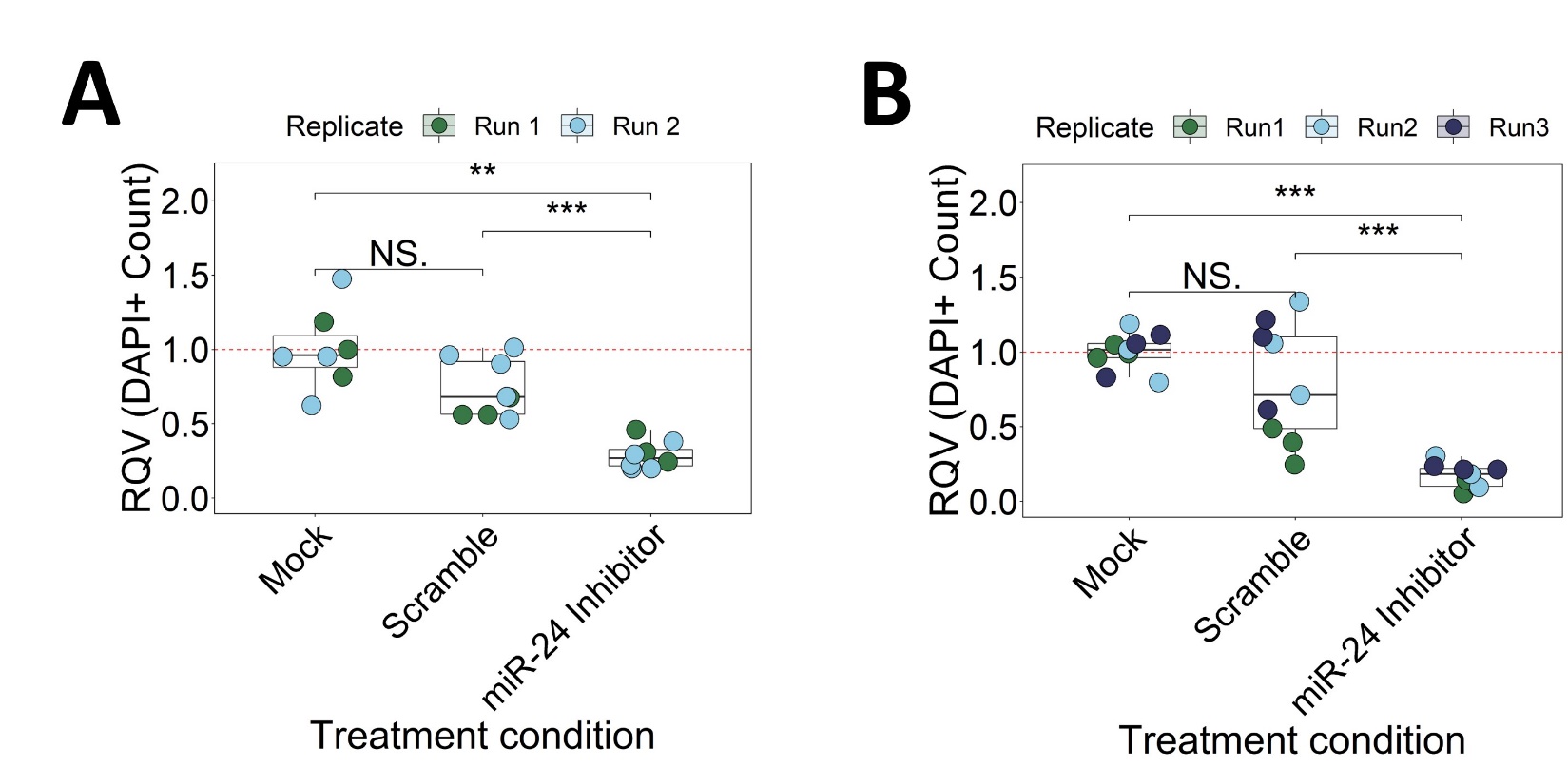

**Supplemental Figure 5:** Relative number of DAPI+ HCT116 cells following mock, scramble or miR-24 inhibitor treatment from (**A**) EdU and (**B**) TUNEL experiments in Figure 5. Significance determined by two-sided Wilcoxon test. Color of data points represents experimental replicate. *p<0.05, **p<0.01, ***p<0.001.

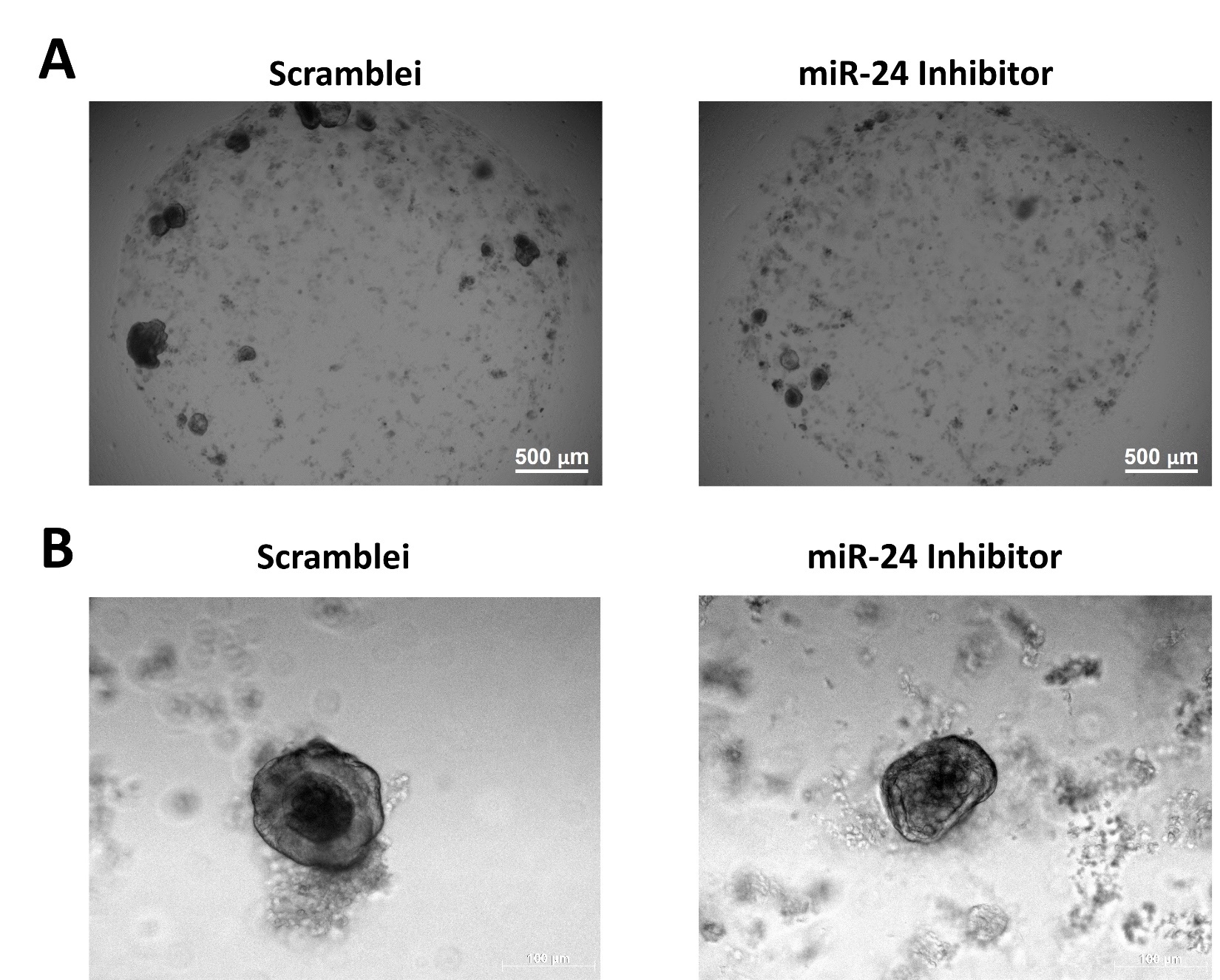

**Supplemental Figure 6:** (**A**) Representative image of WT enteroid wells treated with scramble or miR-24 inhibitor and (**B**) representative image of a single enteroid treated with scramble or miR-24 inhibitor from a second experimental replicate.

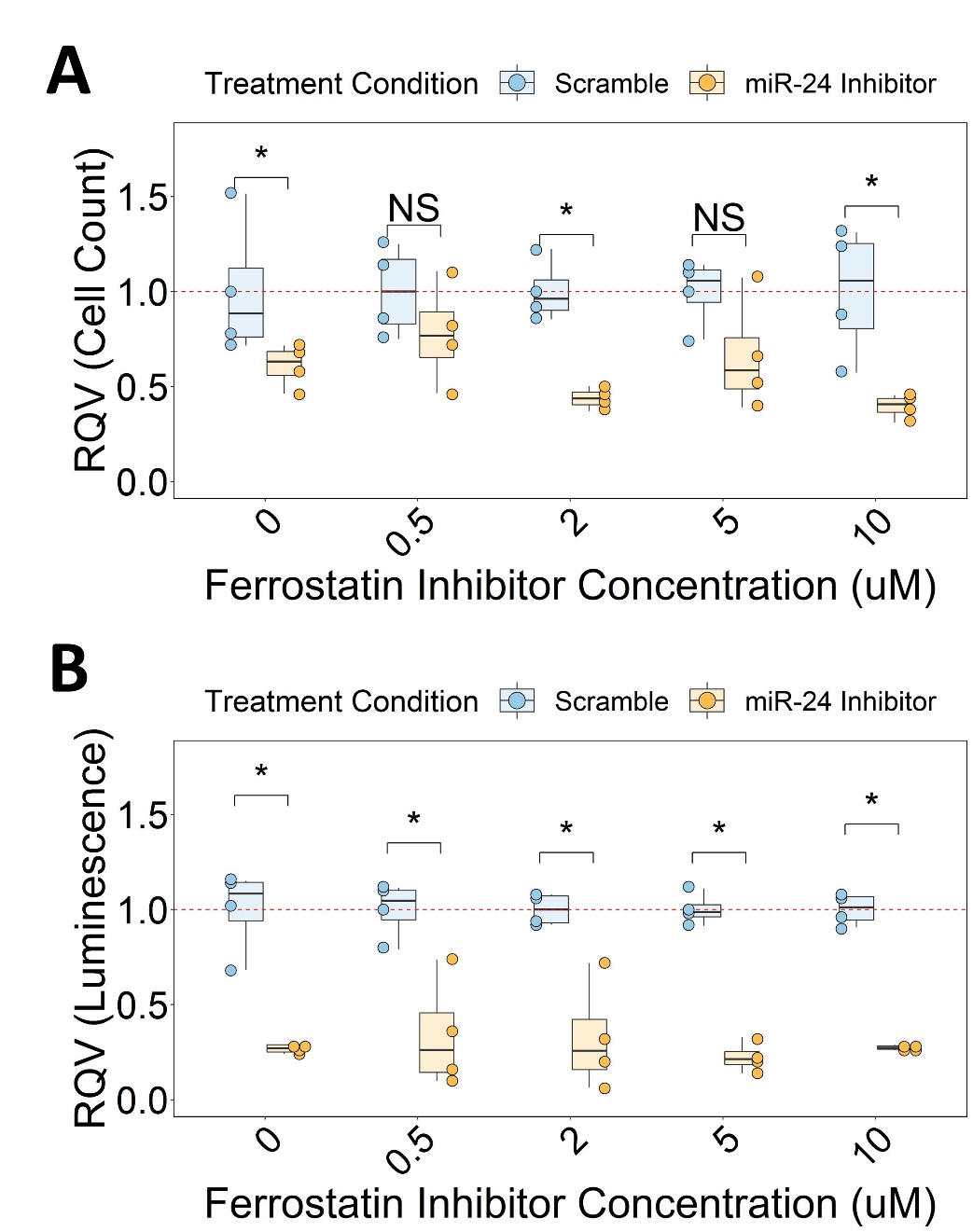

**Supplemental Figure 7:** (**A**) Cell count and (**B**) CellTiter-glo assays following transfection of HCT116 cells with miR-24 inhibitor or scramble along with 0, 0.5, 2, 5, or 10 µM ferrostatin-1. Significance determined by two-sided Wilcoxon test. *p<0.05, **p<0.01, ***p<0.001.
